# Methanol chemoreceptor MtpA- and flagellin protein FliC-dependent methylotaxis determines the spaciotemporal colonization of PPFM in the phyllosphere

**DOI:** 10.1101/2024.08.24.609498

**Authors:** Shiori Katayama, Kosuke Shiraishi, Kanae Kaji, Kazuya Kawabata, Naoki Tamura, Akio Tani, Hiroya Yurimoto, Yasuyoshi Sakai

## Abstract

Pink-pigmented facultative methylotrophs (PPFMs) capable of growth on methanol are dominant and versatile phyllosphere bacteria that provide positive effects on plant growth through symbiosis. However, the spatiotemporal behavior of PPFMs on plant surfaces and its molecular basis are unknown. Here we show that *Methylobacterium* sp. strain OR01 inoculated onto red perilla seeds colonized across the entire plant surface in the phyllosphere concomitant with the plant growth. FliC flagellin proteins required for motility were necessary for distributed colonization on plant leaves, but not for colonization at the leaf periphery. Methanol-sensing chemoreceptor MtpA-dependent chemotaxis (methylotaxis; chemotaxis toward methanol) facilitated the bacterial movement from the peripheral to the inner part of leaves and further entry into the stomatal cavity, indicating that methanol functions as a volatile messenger attracting PPFMs to particular leaf locations possibly to support with photosynthesis and to afford protection from pathogenic invasion.

## Introduction

*Methylobacterium* sp., referred to as pink-pigmented facultative methylotroph (PPFM), is one of the most ubiquitous and abundant bacterial genera present in the aerial parts of the plant, the phyllosphere. PPFM is estimated to exist at 10^4^–10^7^ colony forming units (CFU) per gram fresh weight of plant materials (Delmotte et al., 2009). Many strains of *Methylobacterium* are known for their ability to promote plant growth through the production of plant hormones such as auxins and cytokinin, and to induce systemic resistance to pathogens and diseases (Ardanov et al., 2012; Hornschuh et al., 2006; Koenig et al., 2002; Madhaiyan et al., 2006; Senthilkumar et al., 2009; Yurimoto et al., 2021). In turn, they obtain nutrients from the host plant, establishing the symbiotic relationship in the phyllosphere.

*Methylobacterium* species phylogenetically close to *Methylobacterium* sp. strain OR01 is observed on red perilla seeds and leaves collected from various regions of Japan (Mizuno et al., 2013), indicating that this species is the major colonizer of red perilla leaves. Such species-level specificity of the interaction between the red perilla plant and strain OR01 makes them an ideal system for investigating the ecology and physiology of PPFMs in the phyllosphere. The factors and features that contribute to strain OR01 attaining the dominant colonizer status of perilla phyllosphere still remain unknown and need to be elucidated.

Even though the phyllosphere is speculated to be nutrient-poor, plants indeed provide nutrient compounds sufficient to sustain large microbial communities (Mercier & Lindow, 2000). In particular, methanol, produced from the cell wall component pectin, is recognized as a relatively abundant carbon source for methanol-utilizing microorganisms (Dorokhov et al., 2018), which proves advantageous for PPFMs and allows them to dominate the phyllosphere (Ochsner et al., 2019). PPFMs have developed several survival strategies to help them adapt to the harsh and ever-changing phyllosphere environment, brought about by various factors such as UV, temperature, osmotic and oxidative stresses (Vorholt, 2012). In *Methylorubrum extorquens* AM1, the KaiC protein, a homologue of the cyanobacterial circadian protein, has been shown to facilitate colonization in the phyllosphere through a concerted regulation of environmental responses towards UV and temperature (Iguchi et al., 2018). Previously, we revealed a periodic fluctuation in methanol concentration on *Arabidopsis* leaves, i.e. high in the dark period and low in the light period, using a yeast methanol sensor that could directly measure methanol concentrations (Kawaguchi et al., 2011). We also showed that in response to these diurnal changes in methanol concentration, the methanol-utilizing yeast *Candida boidinii* regulates its gene expression and peroxisome homeostasis necessary for methanol metabolism.

Chemotaxis, the movement of bacteria as a response to chemical stimulus, is driven by three main components: methyl-accepting chemotaxis proteins (MCPs) as sensor chemoreceptors, Che proteins as transmitters of chemotactic signals through phosphorylation, and flagellum as the driving force of directed movement (Hazelbauer et al., 2008; Parkinson et al., 2015) (cf. Fig. 4A). Our recent study on *Methylobacterium aquaticum* strain 22A showed that three MCPs (MtpA, MtpB and MtpC) were responsible for chemotaxis toward methanol, methylotaxis (Tani et al., 2023). A triple *M. aquaticum* strain 22A mutant of these MCPs lost methylotaxis and showed less efficient colonization on plants than the wild-type strain.

In this study, we investigated the factors responsible for the spatiotemporal colonization of PPFM in the phyllosphere. We studied the behavior of *Methylobacterium* sp. strain OR01 at a single-cell level during the growth of red perilla, and followed its entire journey from seeds to the whole plant surface, and subsequently to the next-generation seeds. Fluorescent and electron microscopical observations elucidated the bacterial distribution and dynamic behavior in the phyllosphere. With the analyses of mutants impaired in FliC (the flagellin protein) and MptA of strain OR01, we demonstrate the importance of motility and methylotaxis in phyllosphere colonization of strain OR01 and show a novel function of methanol as a volatile messenger in the phyllosphere during symbiosis between PPFMs and the host plant.

## Results

### Fluorescent single-cell analysis revealed the transmission of PPFM through plant tissue surfaces to seeds for the next generation

*Methylobacterium* sp. strain OR01 expressing GFP (strain OR01-GFP) was inoculated onto seeds of red perilla and the bacterial behavior was followed by fluorescence. Three to four months after inoculation, aerial parts of the plant (the second leaf, newest leaf, petiole, stem, terminal bud, axillary bud and flower, as well as the next-generation seeds picked up from the seed bag of the flower) were harvested at various stages (Fig. 1A). The collected samples were rinsed in sterile water and spread onto a minimal medium (hypho medium) agar plate supplemented with methanol as a single carbon source. GFP-fluorescent colonies appeared on all the tested plates, confirming the presence of strain OR01-GFP on the surface of the entire red perilla plant (Fig. 1B). We also quantified the cell number of the strain OR01 expressing mCherry (strain OR01-mCherry) using flow cytometry (FCM) analysis to understand the localization of the bacteria. Similar to colony formation analysis, aerial parts of the perilla plant were harvested three to four months after strain OR01-mCherry was inoculated onto seeds. FCM analysis revealed that about 13,000 cells were present per mg of the axillary bud, while approximately 2,000 cells were present on the petiole, terminal bud and stem (Fig. 1C). Among the leaf samples, the largest number of cells was detected with new leaf samples. These results indicated that a higher number of bacteria colonize younger plant tissues such as axillary bud and the newest leaves. We also harvested new seeds from flowers and placed them onto a hypho medium agar plate for direct observation by a transilluminator imaging system (Fig. 1D). In addition, cells of strain OR01 were isolated from the seeds placed on the agar plate for microscopic observation (Fig. 1E). We detected GFP fluorescence from the plant and bacterial samples, demonstrating that strain OR01-GFP traveled throughout the surface of plant tissues, and colonized the next-generation seeds.

**Figure 1:**
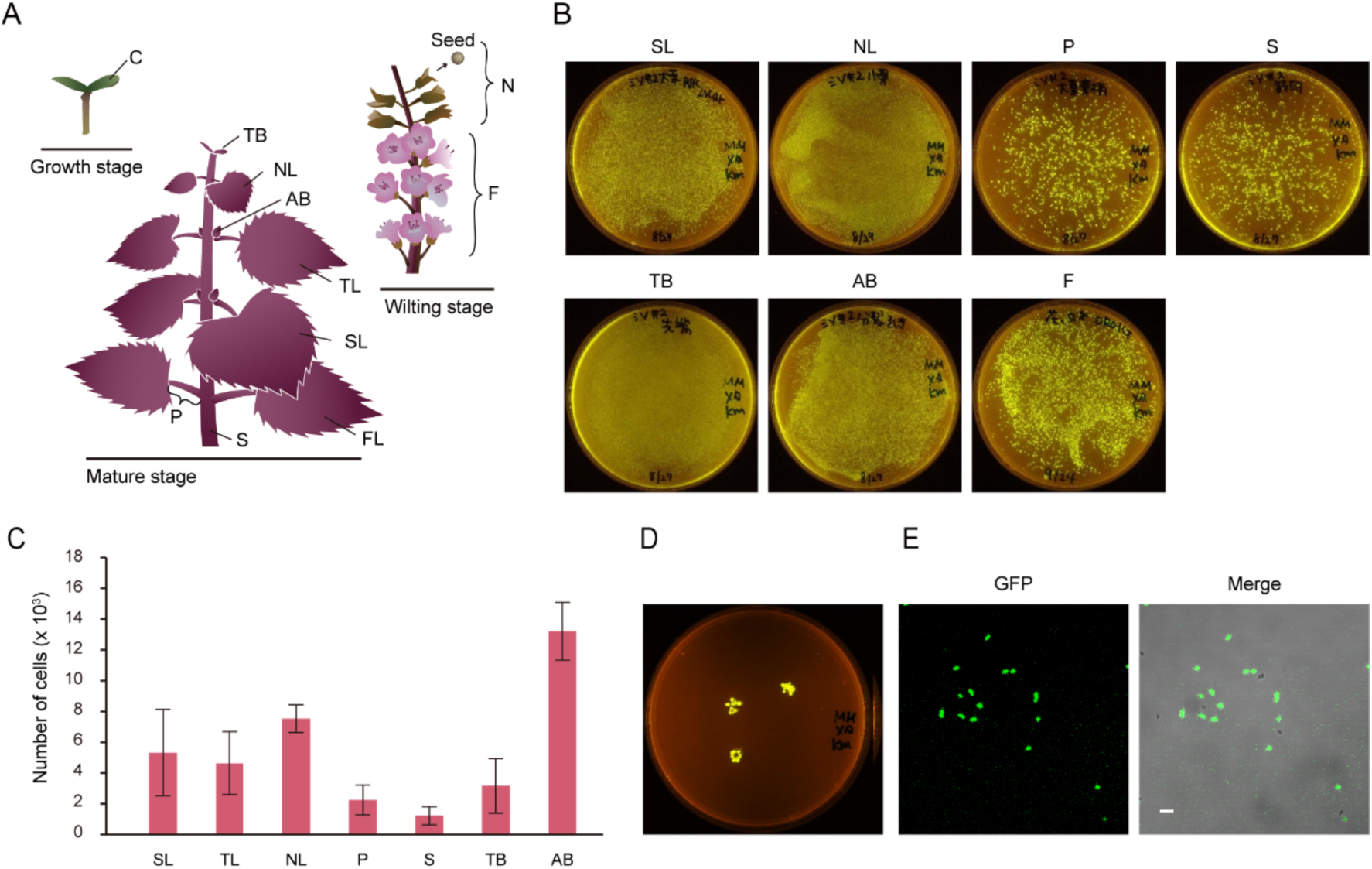
Colonization of *Methylobacterium* sp. strain OR01 from seeds to whole plant of red perilla through plant surface. **A** Schematic images of red perilla during plant development, i.e., growth, mature and wilting stages. C, cotyledon; TB, terminal bud; AB, axillary bud; P, petiole; S, stem; FL, first true leaf; SL, second true leaf; TL, third true leaf; NL, newest leaf; F, flower and N, next-generation seeds. **B** Colony formation activity of the strain OR01-GFP collected from various parts of red perilla. Plant samples were harvested and suspended in sterile water. The bacterial cell suspension was spread onto hypho miminal agar plates supplemented with 0.5% methanol. Strain OR01-GFP was detected by FAS-Digi imaging system after 2–3 days. **C** Quantitation of cell populations of strain OR01-mCherry on red perilla by FCM. Values are indicated as the number of cells per mg of plant sample and are shown as mean ± standard error of the mean (s.e.m.) of at least three independent measurements. **D** The presence of strain OR01-GFP on next-generation seeds. Next-generation seeds were directly placed on hypho minimal agar plates and observed by FAS-Digi imaging system after 3 days. **E** Microscopic images of the strain OR01-GFP collected from next-generation seeds. Cells of strain OR01-GFP were isolated from the seeds used in (**D**) and placed onto a slide glass for observation. GFP-fluorescence image (left) and differential interference contrast (DIC) image were merged (right). Bar, 5 µm.

### PPFM preferentially resides at the base of the trichome, along the vein and around the stomata

To investigate the bacterial distribution on plant leaves, a leaf-print assay was performed with the first true leaves. We found that strain OR01-GFP was distributed throughout both adaxial and abaxial sides of the leaves (Figure 2A). A confocal microscopic analysis found that bacterial colonies were present particularly at the base of the trichomes (Figure 2B left image), along the vein (central image) and around the stomata (right image).

**Figure 2:**
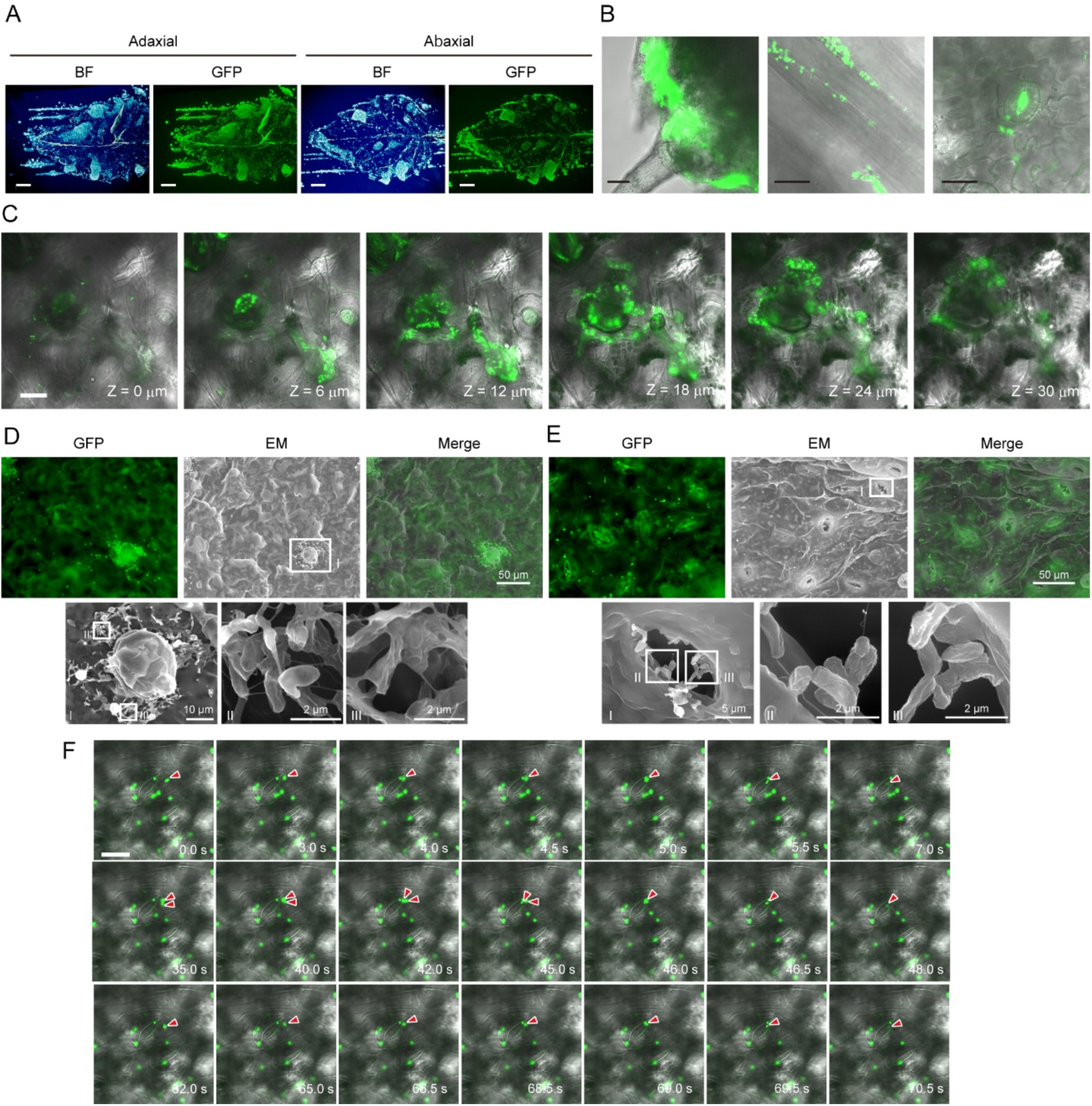
Distribution of *Methylobacterium* sp. strain OR01 on red perilla leaves. **A** Leaf-print assay of strain OR01-GFP on red perilla leaves at the adaxial side (left) and abaxial side (right). Bright-field (BF) image (left) and GFP-fluorescence image (right) are presented. Bar, 2 mm. **B** Confocal fluorescence microscopic images showing strain OR01-GFP distributed at the base of the trichome (left), along the vein (central) and stomata (right). Bar, 20 µm. **C** Z-stack images of stomata with strain OR01-GFP taken by a confocal fluorescence microscope (see also Video 1). Every next image from the left to the right was taken at 6 μm deeper location compared to the previous one. Bar, 20 µm. **D-E** CLEM images of strain OR01-GFP around the trichome (**D**) and stomata (**E**). GFP fluorescence images (upper left) and EM images (upper central) were used for merged CLEM images (upper right). The EM images highlighted with a square (I) were magnified (lower left). The areas marked as square (II) and (III) were further magnified ( (II): lower central and (III) lower right), respectively). Bars show the indicated lengths. **F** Time-lapse images of strain OR01-GFP entering the stomatal cavity. The video was taken by a confocal microscope and recorded for 77 seconds. 21 images extracted at the indicated time points are shown (see also Video 2). Red arrows indicate cells entering into the stomatal cavity from the surface of the leaf. Bar, 20 µm.

Interestingly, GFP fluorescence was detected not only around but also inside the stomata. The z-stack images found that a substantial number of strain OR01-GFP was present in the sub-stomatal cavity (Figure 2C and Video 1).

To further analyze strain OR01-GFP on the leaf surface, correlative light and electron microscopy (CLEM) was performed. We found that numerous clumps were present throughout the leaf surface (Figure 2—figure supplement 1A). Enlarged images revealed that strain OR01-GFP cells were present around the trichome in biofilm-like structures that were about 20 µm x 20 µm long vertically and horizontally (Figure 2D). Furthermore, CLEM captured the presence of strain OR01-GFP around the stomata (Figure 2—figure supplement 1B). Under higher magnification, strain OR01-GFP was detected inside the stomata (Figure 2E), as demonstrated in Figure 2B-C with the confocal microscope. To investigate the bacterial dynamics around the stomata, we monitored the strain OR01-GFP that existed around the stomata over time. Interestingly, the videography captured the moments of strain OR01-GFP entering the stomata (Figure 2F and Video 2). Within the recorded 77 seconds, four cells of strain OR01-GFP moved into the stomata and these cells were observed to track the same path (Video 2). Several non-GFP-labeled cells were also detected, but they did not enter into the stomatal cavity.

### FliC (flagellin)-dependent motility and bacterial distribution in the phyllosphere

Our observation of the cell entry into the stomata prompted us to speculate that bacterial distribution in the phyllosphere is affected by flagellin-dependent motility. We constructed a gene-deletion strain OR01 lacking all of the flagellin proteins, i.e., *fliC1*, *fliC2* and *fliC3*, that expressed GFP (strain Δ*fliC-*triple-GFP). Deletion of these three genes resulted in complete loss of flagella (Figure 3—figure supplement 1A-B) and significant loss of motility (Figure 3—figure supplement 1C-D and Video 3, 4).

We inoculated strain Δ*fliC-*triple-GFP and strain OR01-mCherry on red perilla seeds and harvested the first and second true leaves one month after cultivation. Leaf-print assay revealed that the GFP fluorescence of strain Δ*fliC-*triple-GFP was much weaker than the mCherry fluorescence of strain OR01-mCherry under both single and mixed inoculation conditions (Figure 3A-B). FCM-based quantitative analysis counted approximately 20,000 cells of strain OR01-mCherry and 5500 cells of strain Δ*fliC-*triple-GFP per mg of perilla leaf under single inoculation conditions (Figure 3C). Under mixed inoculation conditions, the difference between these strains became more pronounced in that the cell number of strain OR01-mCherry remained around 15000, whereas that of strain Δ*fliC-* triple-GFP was reduced to less than 300 (Figure 3D). These results suggested that bacterial motility promoted phyllosphere colonization of strain OR01 during transmission from seeds to the phyllosphere. However, strain Δ*fliC-*triple-GFP significantly colonized the peripheral regions around the base of the trichome (Figure 3A-B), indicating that bacterial motility is not responsible for peripheral colonization as much, but is critical for movement toward the center of the leaf surface. This observation could be explained by the assumption that bacteria do not require motility for colony formation in the leaf periphery if they can proliferate on vegetative tissue of the plant (e.g., the meristem of the growing leaf) during the entire growth period.

**Fig 3:**
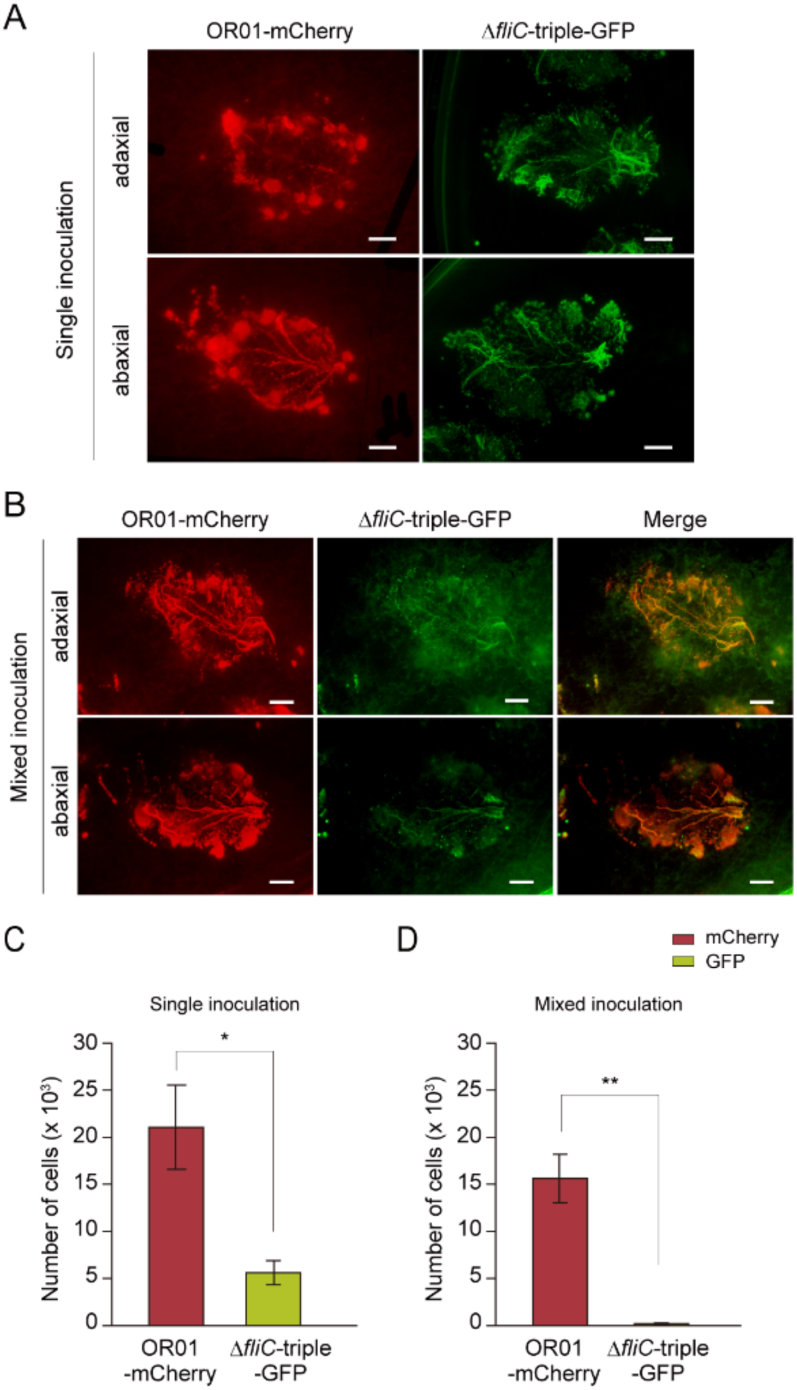
Colonization and distribution of FliC (flagellin)-deleted mutant of strain OR01 in the phyllosphere of red perilla. **A-B** Visualization of colonization and distribution of strain OR01-mCherry and strain Δ*fliC*-triple-GFP on red perilla by leaf-print assay. Strain OR01-mCherry and strain Δ*fliC*-triple-GFP were inoculated in a single inoculum (**A**) or in a mixed inoculum (**B**). Bar, 2 mm. **C-D** FCM-based quantification of cell populations of strain OR01-mCherry and strain Δ*fliC*-triple-GFP on red perilla. Strain OR01-mCherry and strain Δ*fliC*-triple-GFP were inoculated in a single inoculum (**C**) or in a mixed inoculum (**D**). Values are indicated as the number of cells per mg of plant sample and are shown as mean ± s.e.m. of eight independent measurements. Asterisks indicate the level of statistical significance: ** p < 0.01, * p < 0.05.

### MtpA-dependent methylotaxis accelerates stomatal entry and colonization of PPFM to the leaf surface

Bacterial chemotaxis is driven by a signal transduction mechanism involving MCPs and Che proteins and the directional movement controlled by the flagellin protein FliC (Figure 4A). To further investigate the involvement of chemotaxis in phyllosphere colonization, we focused on methylotaxis. We disrupted *mtpA* (homologous to methylotaxis protein A found in *M. aquaticum* strain 22A) (MamtpA, identity: 49%, similarity: 63%) in the genome of strain OR01 (Figure 4—figure supplement 1A-B). Stereo microscopic and FCM analyses using the strain Δ*mtpA* expressing GFP (strain Δ*mtpA*-GFP) revealed that deletion of the *mtpA* gene led to a significant reduction in methylotaxis of strain OR01 without disrupting chemotaxis to succinate (Figure 4—figure supplement 2A-B), indicating that MtpA is a major chemoreceptor for methanol in strain OR01. In a similar manner to strain OR01-GFP, we inoculated strain Δ*mtpA*-GFP onto the abaxial surface and examined the stomatal entry of the mutant strain. Microscopic analysis demonstrated that the cell number of strain Δ*mtpA*-GFP present in the stomata was significantly lesser than that of strain OR01-GFP (Figure 4B). A similar analysis was performed with strain Δ*fliC-*triple-GFP (Figure 4B), which revealed that the lack of FliC proteins resulted in further reduction in the number cells detected compared to that in strain Δ*mtpA*-GFP. Quantitative analysis revealed that 65% of the total area of open stomata was covered by GFP fluorescence of strain OR01-GFP, whereas only 17% and 0.1% were covered by GFP fluorescence of strain Δ*mtpA*-GFP and strain Δ*fliC-*triple-GFP, respectively (Figure 4C).

**Fig 4:**
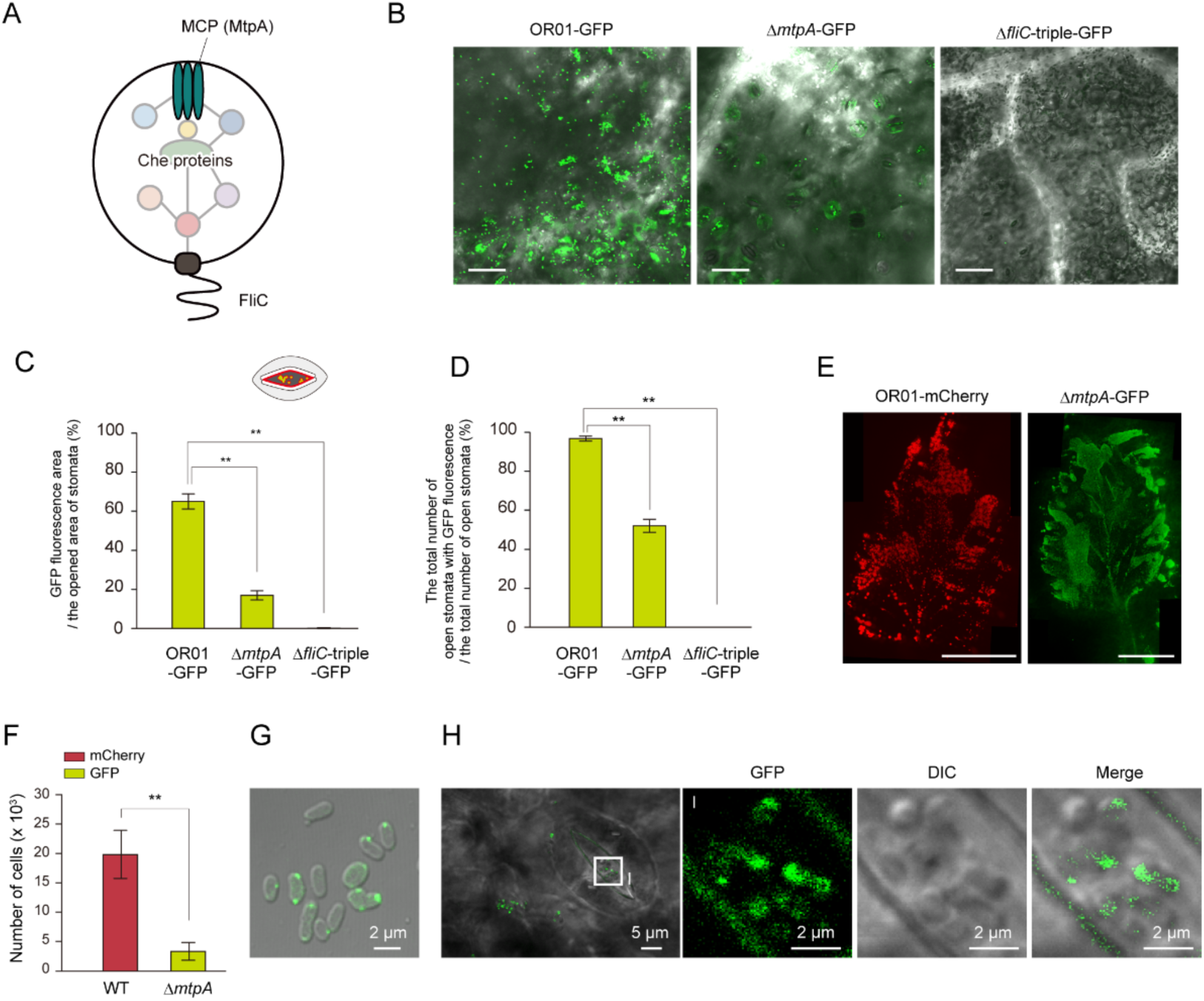
MtpA-dependent methylotaxis affected the distribution and colonization of strain OR01 in the phyllosphere of red perilla. **A** Schematic diagram of the molecular mechanism of methylotaxis, consisting of methanol chemosensor MtpA, Che proteins and flagella (FliC). **B-D** The strain OR01-GFP, strain Δ*mtpA*-GFP and strain Δ*fliC*-triple-GFP present in a single stoma. (**B**) Representative confocal microscopic images of strain OR01-GFP, strain Δ*mtpA*-GFP and strain Δ*fliC*-triple-GFP on red perilla leaves. Marged images of the DIC and GFP images are shown. Bar, 50 µm. (**C**) Quantification of the ratio of (area of the open stomata with GFP fluorescence) / (total area of open stomata) (%). Values are indicated as the number of cells per mg of plant sample and are shown as mean ± s.e.m. of the analysis of 18 stomata. Asterisks indicate the level of statistical significance between strain OR01-GFP and strain Δ*mtpA*-GFP, and strain OR01-mCherry and strain Δ*fliC*-triple-GFP: * p < 0.05, ** p < 0.01. (**D**) Quantification of the ratio of (number of open stomata with GFP fluorescence) / (number of all open stomata) (%). Values are indicated as the number of cells per mg of plant sample and are shown as the mean ± s.e.m. of the analysis of 60 stomata. Asterisks indicate the level of statistical significance between strain OR01-GFP and strain Δ*mtpA*-GFP, and strain OR01-GFP and strain Δ*fliC*-triple-GFP: ** p < 0.01. **E** Leaf-print assay of strain OR01-mCherry and strain Δ*mtpA*-GFP on the second true leaves of red perilla. Strain OR01-mCherry and strain Δ*mtpA*-GFP were inoculated in a single inoculum. Four to six images were combined by ImageJ plug-in MosaicJ for representative images. Bar, 1 cm. **F** FCM-based quantification of cell populations of strain OR01-mCherry and strain Δ*mtpA*-GFP on red perilla. Values are indicated as the number of cells per mg of plant sample and are shown as mean ± s.e.m. of six independent measurements. Asterisks indicate the level of statistical significance between strain OR01-GFP and strain Δ*mtpA*-GFP: ** p < 0.01. **G** Confocal microscopic images of strain Δ*mtpA*-GFP-MtpA. Cells were cultured on hypho minimal medium containing methanol as a carbon source. Marged image of the DIC and GFP images is shown. Bar, 2 µm. **H** Confocal microscopic images of strain Δ*mtpA*-GFP-MtpA on the red perilla leaf surface. The original DIC- and GFP-merged image is shown (left). The highlighted square (I) is magnified, and shown with GFP, DIC, and merged images. Bars show the indicated lengths.

In addition, at least one cell of strain OR01-GFP was observed in more than 95% of the stomata, whereas cells of strain Δ*mtpA*-GFP were present in approximately 52% of the stomata and those of strain Δ*fliC-*triple-GFP were not detected (Figure 4D), demonstrating that MtpA is responsible for strain OR01 to enter into the stomatal cavity. Subsequently, we investigated the physiological significance of MtpA-dependent methylotaxis in bacterial colonization in the phyllosphere of red perilla. After sterilization, red perilla seeds were inoculated with either strain OR01-mCherry or strain Δ*mtpA*-GFP. Two months after plant growth under single inoculation conditions with bacterial cells, we performed the leaf-print assay and found a significant reduction in the fluorescence of GFP from strain Δ*mtpA*-GFP when compared with strain OR01-mCherry (Figure 4E). We also detected a higher proportion of strain Δ*mtpA*-GFP present in the peripheral part of the leaf compared to strain OR01-mCherry. Quantitative analysis by FCM revealed that an average of 20,000 cells per mg of the plant sample was detected from the leaf inoculated with strain OR01-mCherry, whereas only about 4,000 cells of strain Δ*mtpA*-GFP were counted from the collected leaf sample (Figure 4F), suggesting that methylotaxis plays a critical role in the colonization of strain OR01 in the phyllosphere.

Although the *mtpA* deletion led to a significant reduction in the bacterial entry into the stomatal cavity and colonization on the leaf surface, its impact was not as large as the complete loss of flagellin (Figure 4B-D). Next, the chemotactic response to red perilla extract was investigated by fluorescence stereo microscopy and FCM. Strain OR01-mCherry and strain Δ*mtpA*-GFP showed a similar chemotaxis activity to the red perilla extract (Figure 4—figure supplement 2C-D), suggesting that some attractants other than methanol present in perilla extract attract strain OR01, eventually contributing to their stomatal entry and colonization.

To investigate the intracellular localization of MtpA in strain OR01, we constructed the Δ*mtpA* strain expressing GFP-MtpA fusion protein (strain Δ*mtpA*-GFP-MtpA). The functionality of the fusion protein was confirmed by complementation experiment of methylotaxis, using strain OR01, strain Δ*mtpA* and strain Δ*mtpA*-GFP-MtpA (Figure 4— figure supplement 2E). During cultivation on methanol, GFP-MtpA signal was found to form foci and localize mainly to the poles of cells (Figure 4G), suggesting that MtpA in strain OR01 form the chemotaxis sensory array at the cell pole together with Che proteins, in a manner similar to that previously demonstrated in other MCPs (Briegel et al., 2009). Subsequently, we inoculated strain Δ*mtpA*-GFP-MtpA onto the leaf surface of red perilla and found that GFP-MtpA localized at the cell pole (Figure 4H). These results suggested that MtpA is responsible for methylotaxis in the phyllosphere.

### The predominant colonization of strain OR01 on red perilla is attributed to its high methylotaxis activity

Strain OR01 is predominant in the phyllosphere of red perilla (Mizuno et al., 2013). Next, we set out to analyze the reason for this predominance by performing a competitive colonization analysis of mCherry-expressing strain OR01 (strain OR01-mCherry-g) with mVenus-expressing *M. aquaticum* strain 22A (strain 22A-mVenus-g). With a single inoculum of each strain, we found that strain OR01-mCherry-g and strain 22A-mVenus-g were distributed throughout the leaf surface both on the adaxial and abaxial sides (Figure 5A). However, inoculation conditions in a mixed inoculum led to a substantial decrease in the fluorescence of mVenus of strain 22A-mVenus-g cells in comparison with that of strain OR01-mCherry-g on both sides of the leaf surface (Figure 5B). mCherry fluorescence in OR01-mCherry-g was distributed throughout the leaf surface, while mVenus in 22A-mVenus-g was scattered as dots and did not spread uniformly. To quantify the number of cells, the first, second, and third true leaves were collected regularly for FCM analysis. Under inoculation conditions in a single inoculum, ca. 5,000 to 20,000 cells of strain OR01-mCherry-g and ca. 2,500 to 12,000 cells of strain 22A-mVenus-g were detected per mg perilla leaf, and there was no statistical difference between these two strains (Figure 5C). With mixed inoculum of both strains, we found a significant difference in that the cell number of strain OR01-mCherry-g was ca. 5,000 to 25,000, whereas that of strain 22A-mVenus-g was 300 to 3,000 (Figure 5D). These results indicated that strain OR01 is more competitive than strain 22A in colonization on red perilla.

**Fig 5:**
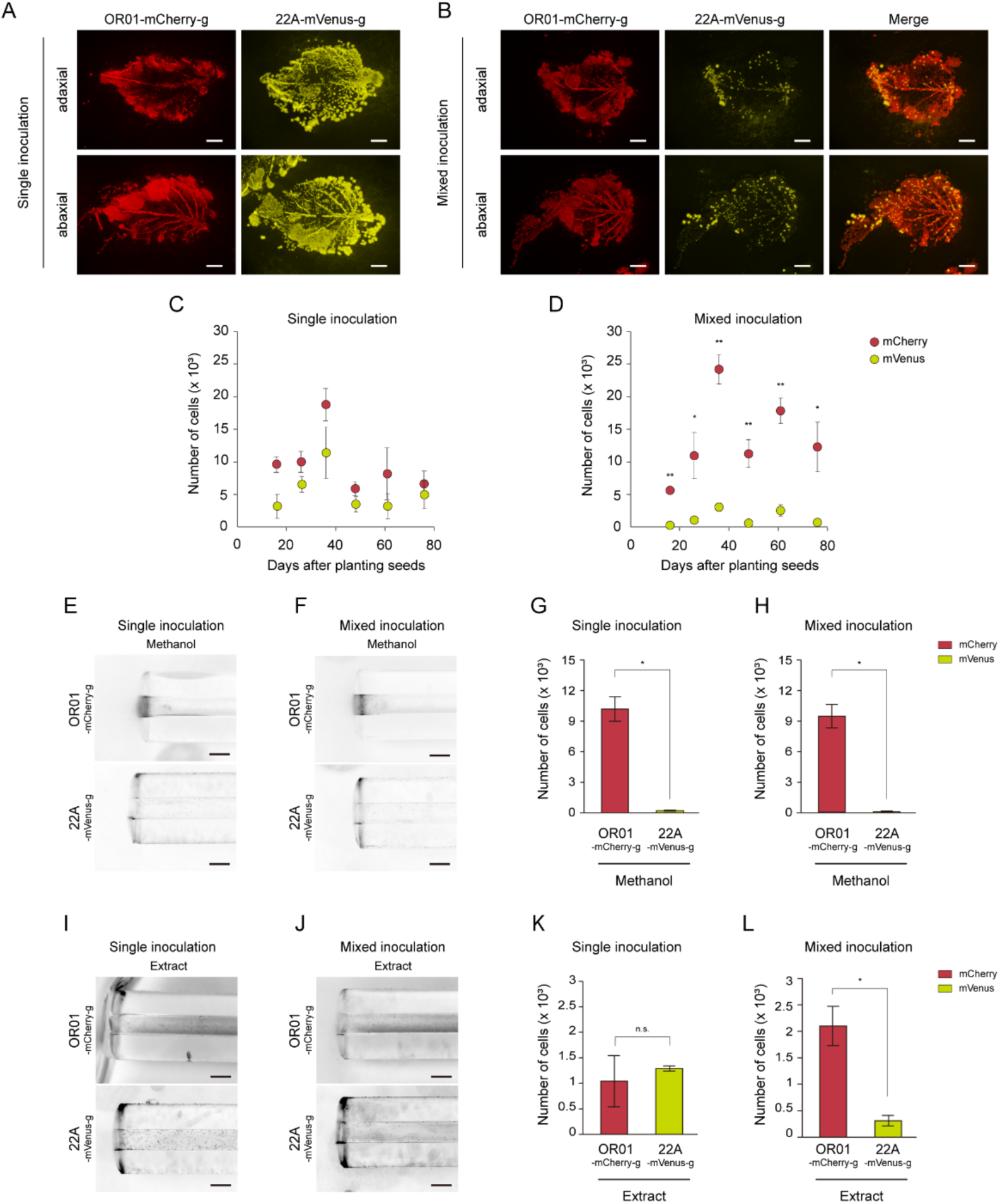
Competitive colonization analysis between strain OR01 and strain 22A on red perilla. **A-B** Leaf-print assay of strain OR01-mCherry-g and strain 22A-mVenus-g on red perilla leaves. These strains were inoculated in a single inoculum (**A**) or a mixed inoculum (**B**). Bar, 2 mm. **C-D** FCM-based quantification of the cell populations of strain OR01-mCherry-g and strain 22A-mVenu-g on red perilla leaves. The first to third true leaves were harvested 16, 26, 36, 48, 61 and 78 days after seed planting with the inoculation of bacterial cells. Strain OR01-mCherry-g and strain 22A-mVenus-g were inoculated in a single inoculum (**C**) or in a mixed inoculum (**D**). Values are indicated as the number of cells per mg of plant sample and are shown as mean ± s.e.m. of at least three independent measurements. Asterisks indicate the level of statistical significance between strain OR01-mCherry-g and strain 22A-mVenu-g on each indicated date: ** p < 0.01, * p < 0.05. **E-H** Capillary assay for methylotaxis of strain OR01-mCherry-g and strain 22A-mVenus-g. (**E-F**) The strains were placed close to the capillaries that contained a hypho minimal medium supplemented with 0.05 % methanol. The strains were inoculated in a single inoculum (**E**) or in a mixed inoculum (**F**). Images were taken by a stereo microscope. Bar, 200 µm. (**G-H**) FCM-based quantification of cell populations of strain OR01-mCherry-g and strain 22A-mVenus-g in capillaries. The strains were inoculated in a single inoculum (**G**) or in a mixed inoculum (**H**). Values are indicated as the number of cells per capillary and are shown as mean ± s.e.m. of three independent measurements. Asterisks indicate level of statistical significance between strain OR01-mCherry-g and strain 22A-mVenus-g: * p < 0.05. **I-L** Capillary assay for chemotactic response of strain OR01-mCherry-g and strain 22A-mVenus-g to red perilla extract. (**I-J**) The strains were placed close to the capillaries that contain a hypho minimal medium supplemented with 0.4% red perilla extract. The strains were inoculated in a single inoculum (**I**) or in a mixed inoculum (**J**). Images were taken by a stereo microscope. Bar, 200 µm. (**K-L**) FCM-based quantification of cell populations of strain OR01-mCherry-g and strain 22A-mVenus-g in capillaries. The strains were inoculated in a single inoculum (**K**) or in a mixed inoculum (**L**). Values are indicated as the number of cells per capillary and are shown as the mean ± s.e.m. of three independent measurements. Asterisks indicate the level of statistical significance between strain OR01-mCherry-g and strain 22A-mVenus-g: * p < 0.05. Not significant: n.s.

Finally, we explored the characteristics that allowed strain OR01 to be competitive on red perilla by a capillary chemotaxis assay. Analysis by stereo microscopy revealed that methylotaxis activity of strain OR01-mCherry-g was much higher than that of strain 22A-mVenus-g (Figure 5E), which became more significant under inoculation conditions in a mixed inoculum (Figure 5F). FCM analysis quantified that the cell number of strain OR01-mCherry-g was approximately 10,000, whereas that of strain 22A-mVenus-g was 200 with a single inoculum (Figure 5G). With a mixed inoculum, the cell number of strain OR01-mCherry-g was 10,000, whereas that of strain 22A-mVenus-g was reduced to 100, which was consistent with the analysis by stereo microscopy (Figure 5H). To explore the sensitivity of strain OR01 to methanol further, we performed a capillary chemotaxis assay supplemented with different concentrations of methanol. We found that strain OR01 migrated into a capillary with the cell numbers showing the correlation with the methanol concentration as low as 0.000012%, demonstrating the remarkable sensitivity of strain OR01 to methanol (Figure 5—figure supplement 1).

In addition, we also performed the capillary chemotaxis assay with the red perilla extract. Analysis of these two strains by stereo microscopy revealed that their chemotactic response was similar to each other under inoculation conditions in a single inoculum (Figure 5I), whereas inoculation conditions in a mixed inoculum resulted in a much higher chemotaxis of strain 22A-mVenus-g to red perilla extract than that of strain OR01-mCherry-g (Figure 5J). FCM analysis quantified with a single inoculum showed that the numbers of cells per µL of red perilla extract were around 1,000 and 1,300 in strain OR01-mCherry-g and strain 22A-mVenus-g with no statistical difference between these strains (Figure 5K). With a mixed inoculum, the cell number of strain OR01-mCherry-g was 2,000, whereas that of strain 22A-mVenus-g was reduced to 300, also consistent with the analysis by stereo microscopy (Figure 5L). These results suggest that methylotaxis activity of strain OR01 significantly enhanced its competitiveness on red perilla compared to other strains on perilla leaves.

## Discussion

This study demonstrates that methanol-sensing MtpA, which drives methylotaxis, determines the spatiotemporal colonization of *Methylobacterium* sp. strain OR01 in the phyllosphere of its natural host, the red perilla plant. During its transmission, strain OR01 was present in the entire leaf surface with a preference to sites around the periphery, vein, trichome and stomata of the leaf surface. Strain OR01 entered the sub-stomatal cavity attracted by methanol and this behavior was attributed to MtpA-driven methylotaxis and FliC-dependent motility. Furthermore, their high methylotaxis activity contributed to the competitiveness of strain OR01 on red perilla (Figure 6).

**Fig 6:**
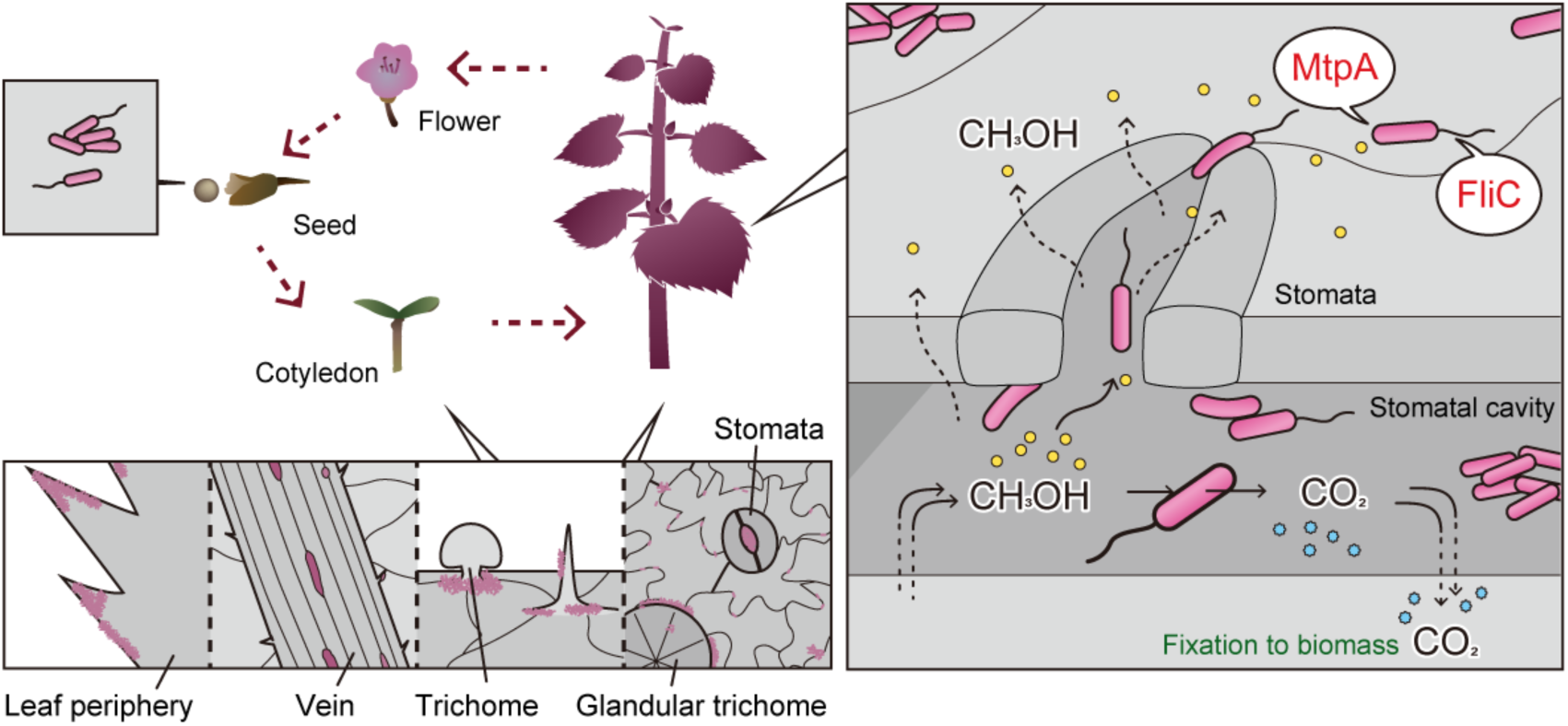
MtpA-dependent methylotaxis and FliC-driven motility enhance the colonization of *Methylobacterium* sp. strain OR-01 in the phyllosphere of red perilla. Strain OR01 travels throughout the plant surface in the phyllosphere and moves to seeds for the next generation concomitant with plant growth. Strain OR01 preferentially colonizes the leaf periphery, veins, trichomes and stomata. Methanol generated from pectin in the plant cell wall functions as a major volatile messenger to attract strain OR01 into the stomatal cavity aided by MtpA-driven methylotaxis and FliC-dependent motility. Within the stomatal cavity, strain OR01 oxidizes methanol to CO_2_, resulting in an increase in CO_2_ concentration used for photosynthesis by the host plant.

Of particular interest is the MtpA-dependent entry of strain OR01 into the stomata (Figure 2F and Video 2). Previously, we developed a methanol cell sensor and directly determined the local methanol concentration on the leaf surface of *Arabidopsis thaliana* to be 0–0.21% (0–64 mM) with a periodic methanol oscillation, i.e. high in the dark and low in the light period (Kawaguchi et al., 2011), which might be directly linked to the hydrolyzation of pectin in the plant cell wall. We confirmed the high methanol-sensing ability of strain OR01 using a capillary assay, which showed that strain OR01 entered the capillary supplemented with 0.000012% methanol (Figure 5—figure supplement 1). The methanol sensing machinery comprising Wsc1 and Wsc3 in the yeast *Komagataella phaffii* senses methanol concentration in the range between 0.005 and 1%. In comparing the bacterial and yeast methanol-sensing machineries, MtpA in strain OR01 is 500-fold more sensitive than the yeast methanol sensor Wsc, suggesting that methanol not only serves as a carbon source for strain OR01, but also functions as a volatile messenger for communication with the host plant.

One possible benefit for the host plant to attract PPFMs to the stomatal cavity is the *in-situ* oxidation of methanol to CO_2_, which is directly fixed by plants through photosynthesis (Figure 6). A previous study with ^13^C-methanol labeling and liquid chromatography-mass spectrometry analyses suggested that methanol present on the leaf surface was primarily oxidized to CO_2_ for energy generation in PPFM, although a portion of the methanol entered into the methanol assimilation pathway (Peyraud et al., 2012). Another benefit of bacterial presence in the stomatal cavity for plants may be the increased protection from pathogenic invasion. *Pseudomonas syringae*, a widely studied plant pathogen, is known to enter the stomatal cavity during infection (Hirano & Upper, 2000; Mansvelt & Hattingh, 1987; Monier & Lindow, 2004; Roos & Hattingh, 1983; Timmer et al., 1987). Therefore, attracted by methanol, PPFMs may move to the stomatal cavity and occupy the colonization place prior to pathogens. Other than the stomatal cavity, invasion of pathogens to the host cells yielded methanol by secreted pectinases (Komarova et al., 2014), which might attract PPFM to the invasion site.

Previously, we reported that many PPFMs isolated from plant samples require B vitamins for growth on minimal media, e.g., pantothenate (vitamin B_5_) (Yoshida et al., 2019). It may be advantageous for PPFMs to colonize sites like glandular trichome to acquire nutrients other than methanol that are necessary for bacterial growth (Tissier et al., 2017).

Deletion of the *mtpA* gene responsible for methylotaxis and *fliC* genes required for motility led to a significant decrease in the number of cells entering the stomata and colonizing red perilla leaves (Figure 4). The role of chemotaxis and/or motility in bacterial colonization has been studied extensively with rhizosphere bacteria, and previous studies have revealed their significance in competitive root surface colonization and establishment of a successful symbiosis (Catlow et al., 1990; Miller et al., 2007; Scharf et al., 2016; Yost et al., 1998). In addition to colonization and symbiosis, transmission is also reported to be supported by bacterial movement (Madsen & Alexander, 1982). The physiological role of chemotaxis has also been investigated using pathogenic bacteria. A previous report has identified AreA and AreB, the genes responsible for chemotaxis in *P. syringae*, as contributing to the pathogenic invasion of the host plant (Ichinose et al., 2023). Another study has found that chemotaxis to nitrate and nitrite contributes to the colonization of the pathogen *Dickeya dadantii* on its host plant (Gálvez-Roldán et al., 2022). However, insights from these studies have been limited to the rhizosphere, in particular to nitrogen fixation bacteria and the loci for pathogenic invasion. Our study has shed light on bacterial chemotaxis and motility of phyllosphere microorganisms by demonstrating their crucial role in colonizing the host plant through its entire life cycle, rather than being confined to a specific part or developmental stage of the plant. To our knowledge, this is also the first report to show methanol as a compound that attracts phyllosphere bacteria to enter stomatal pores.

## Materials and Methods

### Bacterial strains and culture conditions

The bacterial strains used in this study are listed in Supplementary File 1A. *Methylobacterium* sp. strain OR01 and *M. aquaticum* strain 22A were grown at 28°C in the hypho minimal medium (Supplementary File 1B) (Yost et al., 1998) containing one of the carbon sources described below and all of the B-group vitamins (Vitamin mix). Carbon sources were 0.5% or 0.05% (v/v) methanol, 0.5% or 0.05% (w/v) disodium succinate or 0.4% (w/v) extract from red perilla leaves. The antibiotic kanamycin (Km, 20 µg/mL) was added as needed. *Escherichia coli* HST08 premium competent cells (TaKaRa) were used for gene cloning. *E. coli* was grown at 37°C on LB medium (1% Bacto tryptone, 0.5% Bacto yeast extract and 0.5% NaCl) in the presence of ampicillin (50 µg/mL) or kanamycin (20 µg/mL). The nucleotide sequences of *fliC1*, *fliC2*, *fliC3*, *mtpA* and *mxaF* were deposited in the DDBJ/EMBL/GenBank under accession numbers LC833873, LC833874, LC833875, LC833876 and LC833877, respectively.

### Construction of plasmids

The plasmids used in this study are listed in Supplementary File 1C, and the oligonucleotide primers are listed in Supplementary File 1D. The plasmid vectors pBluescript II SK (+) (Stratagene Inc), pDCG-1 (Iguchi et al., 2013), pMO149 (Maeda et al., 2015), pAT02-V (Tani et al., 2023), pCM80-Km (Orita et al., 2014), pCM1682 (Iguchi et al., 2018) and pK18 mobsacB (Iguchi et al., 2013) were used in previous studies.

The vector pBS-P_mxaF_ was constructed as follows: The promoter region of the gene *mxaF* was PCR-amplified by using the genomic DNA of *Methylobacterium* sp. strain OR01 with a primer pair PmxaF-Fw-KpnI/PmxaF-Rv-HindIII. After digestion with *Kpn*I and *Hind*III, the DNA fragment was ligated with linearized pBluescript II SK (+) with the same set of restriction enzymes. Subsequently, the primer pairs GFP-Fw-HindIII/GFP-Rv-PstI and mCherry-Fw-HindIII/mCherry-Rv-BamHI were used to amplify the 0.7-kb *GFP* and *mCherry* genes with pDCG-1 and pMO149 as templates, respectively. The PCR products with *GFP* and *mCherry* sequences were ligated into the linearized pBS-PmxaF by restriction enzyme sets, *Hind*III-*Pst*I and *Hind*III-*Bam*HI, respectively, resulting in the pBS-PmxaF-GFP and pBS-PmxaF-mCherry vectors.

The pCM802-based vectors were constructed from a plasmid vector pCM80-Km (Orita et al., 2014) given by Dr Toshiaki Fukui. The *mxaF* promoter region of *M. extorquens* strain AM1 was removed using a primer set pCM80km-Re1800/pCM80km-Fw2471 by inverse PCR with pCM80-Km as a template, followed by self-ligation, yielding pCM802-Km. The vectors pCM802-Km and P_mxaF_-GFP were digested by *Kpn*I and *Pst*I and ligated to each other, resulting in the plasmid pCM802-GFP. Similarly, the vectors pCM802-Km and P_mxaF_-mCherry were cut by *Kpn*I and *Sph*I, and these DNA fragments were ligated to obtain the plasmid pCM802-mCherry. The vector pCM802-GFP-mtpA was constructed as follows: The promoter and ORF regions of *mtpA* were PCR-amplified using primer pairs NB_PmtpA_Fw/NB_PmtpA+GFP_Rv and NB_PmtpA+GFP_Fw/NB_GFP+mtpA_Rv with the genomic DNA of strain OR01 as templates. The *GFP* coding sequence was amplified by PCR using a primer pair NB_GFP+mtpA_Fw/NB_mtpA_Rv with pCM802-GFP as a template. These two DNA fragments were cloned into the *Bam*HI and *Sac*I sites of pCM802-Km by NEbuilder cloning kit (New England Biolabs), resulting in the vector pCM802-GFP-mtpA. These pCM802-based vectors were used as vectors for expression in plasmids.

The plasmid vectors pCM1684 and pCM1685 for genome insertion into strains OR01 and 22A, respectively, were constructed as follows: pCM1684 was constructed by ligating pCM1682-derived sequences with a strain OR01 chromosome region at 404241 locus where no gene was assigned, and further with P_mxaF_-mCherry sequence amplified from pCM802-P_mxaF_-mCherry. Similarly, pCM1685 was constructed by ligating pCM1682-derived sequences with a strain 22A chromosome region at 1718735 locus where no gene was assigned, and further with P_mxaF_-Venus sequence amplified from pAT02-Venus. These ligation processes were achieved by an NEbuilder cloning kit, resulting in the vector pCM802-GFP-mtpA.

The plasmid vectors for precise in-frame deletion of *mtpA*, *fliC1*, *fliC2* and *fliC3* genes were constructed as follows: pK18mobsacB carrying a positive selection marker (Km^r^ gene) and a counter-selectable marker (suicide gene *sacB*) was used for constructing all the pK18-based plasmid vectors for gene disruption, as previously described^30,31^. Subsequently, the disruption vector pK18 ΔfliC1 was constructed as follows: The *fliC1* and the homologous fragment flanking *fliC1* were PCR-amplified using a primer pair fliC1_up_Fw/ fliC1_down_Rv with the genomic DNA of strain OR01 as a template. The DNA fragment and pK18mobsacB, digested by restriction enzymes *Sal*I and *Sph*I, were ligated to each other, resulting in the plasmid pK18 fliC1. The *fliC1* ORF region was removed using a primer set Inv_fliC1_Fw1/ Inv_fliC1_Rv by inverse PCR with pK18 fliC1 as a template, followed by a self-ligation, yielding the vector pK18 ΔfliC1. The disruption vectors pK18 ΔfliC2 and pK18 ΔmtpA were constructed in a similar manner. The ORF, upstream and downstream region of *fliC2* and *mtpA* were PCR-amplified using primer pairs fliC2_up_Fw/ fliC2_down_Rv and mtpA_up_Fw/mtpA_down_Rv, respectively, with the genomic DNA of strain OR01 as a template. The DNA fragments and pK18mobsacB, digested by restriction enzymes *Sal*I and *Sph*I, were ligated to each other, resulting in the plasmid pK18 fliC2 and pK18 ΔmtpA. The disruption vector pK18 ΔfliC3 was constructed as follows: the homologous fragments flanking *fliC3* were amplified by PCR using primer pairs fliC3_up_Fw/ fliC3_up_Rv and fliC3_down_Fw/ fliC3_down_Rv with the genomic DNA of strain OR01 as a template. These fragments were cloned into the *Sal*I and *Sph*I sites of pK18mobsacB by an NEbuilder cloning kit.

### Transformation of DNA to *Methylobacterium* sp. OR01 and *M. aquaticum* strain 22A

Cells were grown for 48 hours and 500 µL of them were centrifuged at 15,000 rpm for 1 minute. To prepare for competent cells of *Methylobacterium* sp. OR01, harvested cells were washed with sterilized water and then resuspended in 50 µL of sterilized water. For *M. aquaticum* strain 22A, collected cells were washed with 10% glycerol and then resuspended in 50 µL of sterilized water containing 10% glycerol and 30% PEG. Competent cells were mixed with 1 µL of DNA and transferred to 1 mm gap cuvette (Bio-Rad). The DNA was introduced to the cells by electroporation at 1.8–2 kV, 25 µF, 200 Ω (GenePulser Xcell^TM^, Bio-Rad). The entire electroporated suspension was inoculated into 500 µL Nutrient Broth (Difco^TM^) medium and cultured for 3 hours at 28°C. These cells were plated on the hypho medium plates containing 0.5% succinate and vitamin mix and incubated at 28°C for 3–5 days. For obtaining gene knockout strains, the grown colonies were streaked on R2A agar medium containing 10% sucrose, 0.5% succinate and 1% vitamin mix. After culturing at 28°C for about 5 days, mutant strains were selected by colony PCR.

### Cultivation of red perilla

Red perilla was grown in a plastic cup about 20 cm high with vermiculite and a ventilated seal or in a plastic dish about 4 cm high with Hoagland agar as described previously (Mizuno et al., 2013). The plants were placed in the NK Biotron LH-220 (Nippon Medical and Chemical Instruments, Osaka, Japan). The system was operated at 25°C, 65% humidity under a 15-h light 9-h dark cycle. Plants cultivated on agar media developed in the same way as plants grown in plastic cups containing vermiculite for the first 2 weeks. Thereafter, a dwarf phenotype was observed, but they appeared healthy otherwise.

### Measurement of cell populations of PPFMs on red perilla

Cells of the strain OR01 and strain 22A were grown on the hypho medium containing 0.5% methanol and vitamin mix at 28°C for 2 days. After collection, cells were washed with sterilized water and suspended in sterilized water to obtain a suspension with an OD_600_ of 0.1. Red perilla seeds were treated with 70% ethanol for 1 minute and with 1% (v/v) antiformin (containing 0.3% v/v Tween 20) for 5 minutes, followed by washing with sterilized water 5 times. They were soaked in 1 mL of the single or mixed (wild-type strain and each mutant strain) cell suspension for 3 hours with gentle shaking at 5 rpm using a Rotator RT-5 (Taitec, Saitama, Japan) at 28°C. The seeds incubated with *Methylobacterium* sp. OR01 were sown onto Hoagland agar in a plant culture dish (100×40 mm) or vermiculite for growth in the chamber. For the collection of PPFMs from red perilla, the leaves were cut and placed in a 1.5 mL tube, weighed, and then,100 µL of PBS was added per 10 mg of leaf and vortexed for 15 minutes. 20 µL of this collection sample was mixed with 20 µL of BD^TM^ Liquid counting Beads (BD Biosciences) and 200 µL of PBS and the total volume was used for FCM, which was performed as previously described (Tani et al., 2023).

### Colony formation analysis

After sterilization, as previously described (Mizuno et al., 2013), red perilla seeds were suspended in cells of strain OR01-GFP for 3 hours and sown onto Hoagland agar or vermiculite. Three to four months after aseptic growth, aerial parts of the plant were harvested. The collected samples were rinsed, suspended in sterile water and spread onto the hypho medium agar plate containing 0.5% methanol with kanamycin (20 µg/mL) as needed. To visualize colonies, the FAS-Digi imaging system (NIPPON Genetics) was employed.

### Microscopic observations

Fluorescence stereo microscope SZX16 (OLYMPUS), confocal microscopes Zeiss LSM510 META/Axiovert 200 (Olympus), FV3000 (Olympus), and FV4000 (Olympus), BX51 (Olympus), SU8220 (Hitachi High-Tech) and IX73 inverted microscope (Olympus) were used in this study. The images were captured and processed using Olympus cellSens software. Details of these microscopes are as follows:

Fluorescence stereo microscope SZX16 is equipped with a digital charge-coupled device camera (Olympus DP80), a GFP filter (Olympus SZX2-FGFPHQ) and an RFP filter (Olympus SZX2-FRFP2). SZX16 was used for Figure 2A, 3A, 3B, 4E, 5A, 5B, 5E, 5F, 5I, 5J, and Figure4—figure supplement 1A and 1C.

Zeiss LSM510 META/Axiovert 200 is equipped with a Plan Fluor 100×/1.45 NA oil objective. GFP and mVenus fluorescence were obtained with a multiline 488 argon laser and a 505–550 nm filter for emission. An HFT 405/514 beam splitter was used as a connecting filter. Zeiss LSM510 META/Axiovert 200 was used for Figure 1E, 2B, 2C, 2F and Video 1 and 2.

FV3000 is equipped with six solid-state diode lasers (405 nm, 445 nm, 488 nm, 514 nm, 561 nm, and 640 nm), whereas FV4000 comes with ten laser lines ranging from 405 nm to 785 nm. They use Olympus objectives including PLAPON 60x (1.42 NA, oil-dipped) and UPLSAPO 100x (1.35 NA, Si oil-dipped). For image detection, the FV3000 is equipped with high-sensitivity GaAsP detectors, enabling the detection of faint fluorescence signals with high signal-to-noise ratios. FV4000 uses the SilVIR detector, which combines a silicon photomultiplier (SiPM) with advanced signal processing. FV3000 was used for Figure 4B and 4G, while FV4000 was used for Figure 4H.

BX51 is equipped with a digital charge-coupled device camera (Olympus DP71). It uses Olympus objectives including UPlanSApo 4x/0.16, UPlanSApo 10x/0.40, UPlanSApo 20x/0.75 and UPlanSApo x40/0.95. The mirror unit for GFP is U-MNIBA3 (Olympus). BX51 was used for Figure 2D, 2E and Figure 2—figure supplement 1.

SU8220 was used at 15 or 20 kV accelerating voltage. The scanning mode for high magnification is SE (UL). The electron microscope was used for Figure 2D, 2E and Figure 2—figure supplement 1.

IX73 inverted microscope is equipped with a lens UPLAPO OI3 100x / 1.35 and flagellar length was measured by CellSens using the data taken as differential interference contrast (DIC) images. The microscope was used for Figure3—figure supplement 2A, 2C, 2D, Video 3 and 4.

### Capillary chemotaxis assay

Chemotaxis was evaluated using capillaries and a glass slide with a hole^27^. Two capillaries were placed on each side of the hole as spacers between the cover glass and the glass slide. Hypho medium supplemented with a carbon source at a concentration of 0.05%, as necessary, vitamin mix and appropriate antibiotics were sucked into a new capillary. Then, one hole was closed with nail polish and placed on the hole glass slide, followed by a cover glass set over the capillary. For red perilla extract, 1 mL of hypho medium was added to 4.1 mg of red perilla leaves and mashed with a mortar and pestle. Then, the obtained extracts were filtered for experiments. 200 µL of bacterial suspension of OD_600_ at 0.01 was applied to the hole portion of the glass slide with a hole and incubated at 28°C for 3 hours. After culturing, the cells in the capillary were observed with an SZX16 fluorescence stereo microscope (OLYMPUS). The number of cells in the capillary was examined as previously described (Tani et al., 2023).

### Microscopic analysis of cell entry to a single pore of the stomata

Leaves were placed statically overnight on a bacterial solution containing hypo medium with no carbon source. The bacterial solution was prepared at OD_600_ 0.1. The leaves were then observed by confocal microscopes FV3000 (Olympus) and FV4000 (Olympus) for quantitative analysis.

### Leaf-print assay

The harvested leaves of red perilla were cut and placed directly on the hypho medium agar plate containing 0.5% methanol or 0.5% succinate with the addition of kanamycin (20 µg/mL), as needed. Then, the leaf was gently pressed to the medium using a spreading stick and incubated at 28°C for 3–4 days. After the culture, microbial distribution was observed by the SZX16 fluorescence stereo microscope (OLYMPUS).

### CLEM analysis

Red perilla seeds were sterilized and treated with the cell suspension of strain OR01-GFP. They were sown onto Hoagland agar in a plant culture dish for growth in a chamber. One month after aseptic growth, aerial parts of the plant were harvested. Samples were fixed with a solution of 2% paraformaldehyde, 2.5% glutaraldehyde, 0.1 M phosphate buffer, pH 7.4 for 30 minutes at room temperature. Subsequently, these samples were used for CLEM analysis. After fixation, these samples were suspended and stored in phosphate-buffered saline. Subsequently, the samples were mounted on a glass slide and fluorescent and Bright-field images were acquired with BX51. After washing in 0.1 M phosphate buffer (pH7.4) three times, the samples were frozen in liquid nitrogen for 2 minutes and freeze-dried overnight (Yamato Scientific Co., ltd. DC-56A). Then, the samples were mounted on SEM stubs using double-sided carbon tape and coated with 7 nm osmium in an HPC-1SW coater (Shinku Device). A scanning electron microscope (Hitachi High-Tech SU8220) was used for imaging according to the manufacturer’s protocol. After image acquisition, images were merged using Photoshop 2024 (Adobe).

### Flagella visualization

Living cells were stained for flagella visualization using the staining solution by Ryu (Ryu, 1937) and a method similar to Heimbrook and colleagues (Madhaiyan et al., 2006). Visualized flagella were investigated with an IX73 inverted microscope (Olympus) and the length of flagella was analyzed by cellSens.

## Supplementary File information

Supplementary File 1A-D

## Author details

**Shiori Katayama**

Graduate School of Agriculture, Kyoto University, Kitashirakawa-Oiwake, Sakyo-ku, Kyoto, Japan

**Contribution**: Conceptualization, Investigation, Formal analysis, Visualisation

**Competing interests**: No competing interests declared

**Kosuke Shiraishi**

Graduate School of Agriculture, Kyoto University, Kitashirakawa-Oiwake, Sakyo-ku, Kyoto, Japan

**Contribution**: Conceptualization, Formal analysis, Writing – original draft preparation, Funding acquisition

**Competing interests**: No competing interests declared

**Kanae Kaji**

Graduate School of Agriculture, Kyoto University, Kitashirakawa-Oiwake, Sakyo-ku, Kyoto, Japan

**Contribution**: Investigation, Formal analysis

**Competing interests**: No competing interests declared

**Kazuya Kawabata**

Graduate School of Agriculture, Kyoto University, Kitashirakawa-Oiwake, Sakyo-ku, Kyoto, Japan

**Contribution**: Investigation, Formal analysis

**Competing interests**: No competing interests declared

**Naoki Tamura**

Department of Anatomy and Histology, Fukushima Medical University School of Medicine, Hikarigaoka, Fukushima, Japan

**Contribution**: Investigation

**Competing interests**: No competing interests declared

**Akio Tani**

Institute of Plant Science and Resources, Okayama University, Okayama, Japan **Contribution**: Conceptualization, Investigation, Funding acquisition **Competing interests**: No competing interests declared

**Hiroya Yurimoto**

Graduate School of Agriculture, Kyoto University, Kitashirakawa-Oiwake, Sakyo-ku, Kyoto, Japan

**Contribution**: Conceptualization, Formal analysis, Writing – review & editing, Supervision, Funding acquisition

**Competing interests**: No competing interests declared

**Yasuyoshi Sakai**

Graduate School of Agriculture, Kyoto University, Kitashirakawa-Oiwake, Sakyo-ku, Kyoto, Japan

**Contribution**: Conceptualization, Formal analysis, Writing – review & editing, Supervision, Funding acquisition

**Competing interests**: No competing interests declared

## Funding

**JST GteX Program (JPMJGX23B4)**

- Kosuke Shiraishi

**Joint Usage/Research Center, Institute of Plant Science and Resources, Okayama University**

- Akio Tani

**JSPS KAKENHI (19K22307)**

- Hiroya Yurimoto

**JSPS KAKENHI (22K19133)**

- Hiroya Yurimoto

**JSPS KAKENHI (24H02120)**

- Hiroya Yurimoto

**JSPS KAKENHI (19H02870)**

- Yasuyoshi Sakai

## Acknowledgments

We thank Dr Toshiaki Fukui and Dr Izumi Orita for gifting us the plasmid vector pCM80-Km.

---

**Figure 2—figure supplement 1.**
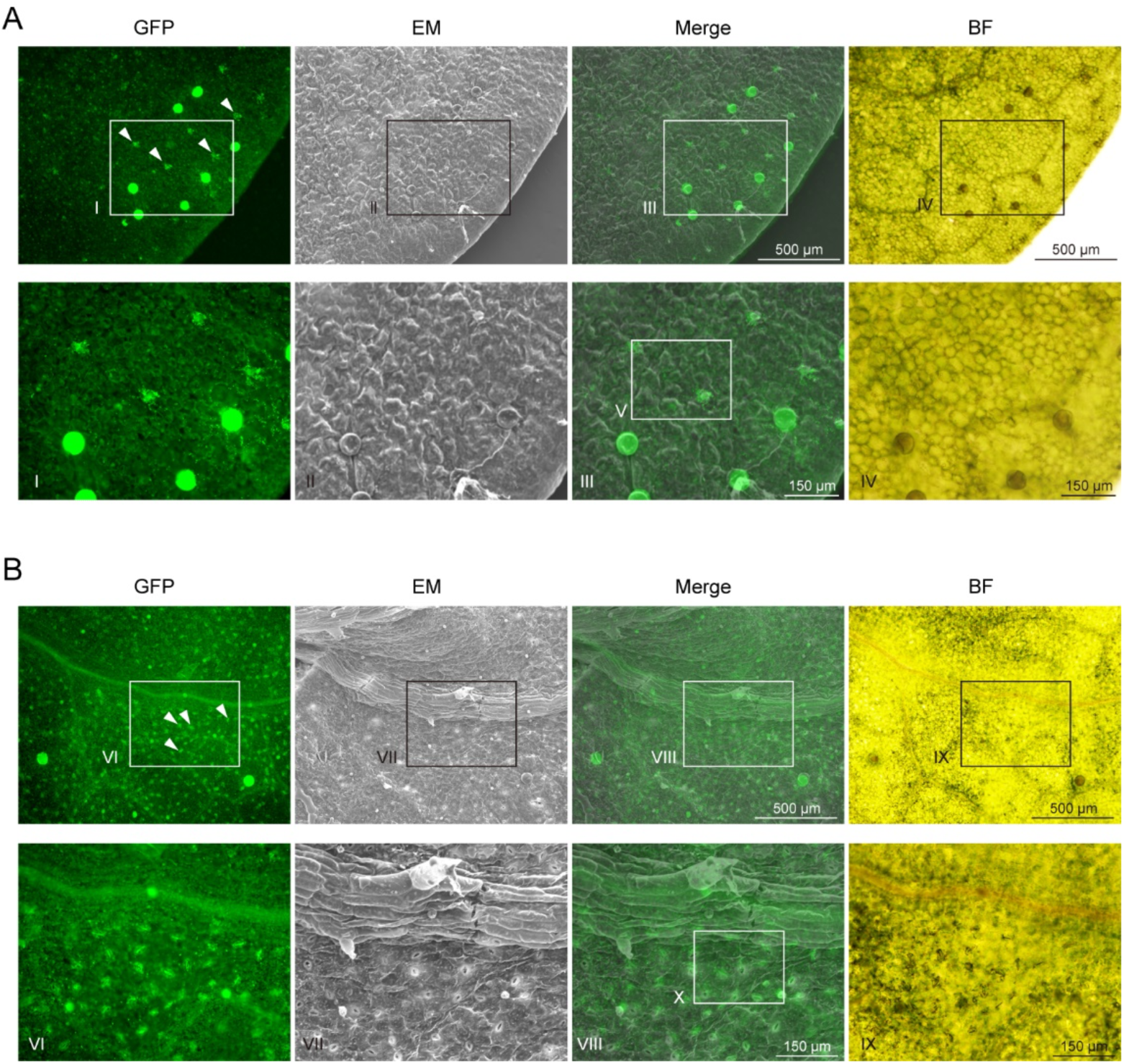
CLEM analysis of strain OR01-GFP colonizing red perilla leaves. **A** CLEM images of strain OR01-GFP at a low magnification with a focus on the trichome (upper panels). GFP fluorescence images (left) and EM images (second from the left) were used for merged CLEM images (second from the right). Bright-field (BF) image is also shown (right). The images highlighted with squares (I), (II), (III) and (IV) were magnified (lower panels). The highlighted square (V) was the merged image used for Fig. 2D (upper panel). White arrows in the GFP image are clumps of strain OR01-GFP. Bars show the indicated lengths. **B** CLEM images of strain OR01-GFP at a low magnification with a focus on the stomata (upper panels). GFP fluorescence images (left) and EM images (second from the left) were used for merged CLEM images (second from the right). Bright-field (BF) image is also displayed (right). The images highlighted with squares (VI), (VII), (VIII) and (IX) were magnified (lower panels). The highlighted square (X) was the merged image used for Fig. 2E (upper panel). White arrows in the GFP image shows bacteria colonizing around the stomata. Bars show the indicated lengths.

**Fig 3—figure supplement 1.**
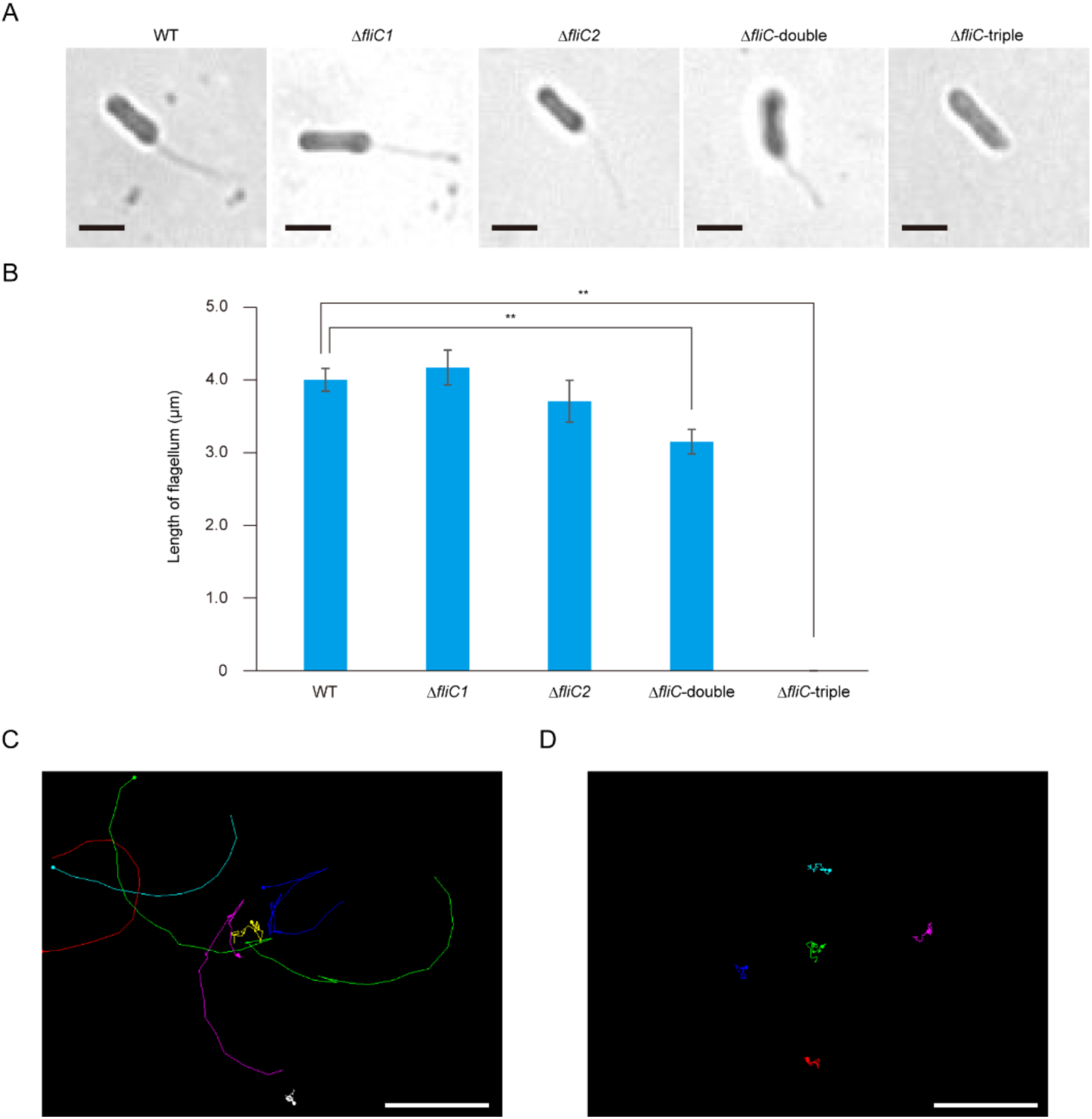
Deletion of *fliC* genes led to the loss of flagella. **A** Inverted microscopic images of wild-type (WT), Δ*fliC1*, Δ*fliC2*, Δ*fliC1*Δ*fliC2* (Δ*fliC*-double) and Δ*fliC1*Δ*fliC2*Δ*fliC3* (Δ*fliC*-triple) cells of strain OR01. Flagella were visualized with *Ryu* stain, but could not be observed in strain Δ*fliC*-triple. Bar, 2 µm. **B** Quantitation of the flagellum length measured from images taken by inverted microscope shown in (**A**). Values are indicated as the number of cells per mg of plant sample and are shown as the mean ± s.e.m. of a minimum of 11cells analyzed. Asterisks indicate the level of statistical significance between wild-type (WT) cells and Δ*fliC*-double cells, and WT cells and Δ*fliC*-triple cells: ** p < 0.01. **C-D** The representative image plots of cell movement. (**C**) Single-cell tracking was analyzed by Manual Tracking plug-in for ImageJ. Plots were created with WT cells during the microscopic observation for 5.8 seconds. The period from 0.68 seconds to 6.48 seconds in Video 3 is shown in this figure. Seven cells were monitored. (**D**) Plots were created with Δ*fliC*-triple cells during the microscopic observation for 5.8 seconds. The period from 0.68 seconds to 6.48 seconds in Video 4 is shown in this figure. Five cells were monitored. Bar, 20 µm.

**Fig 4—figure supplement 1.**
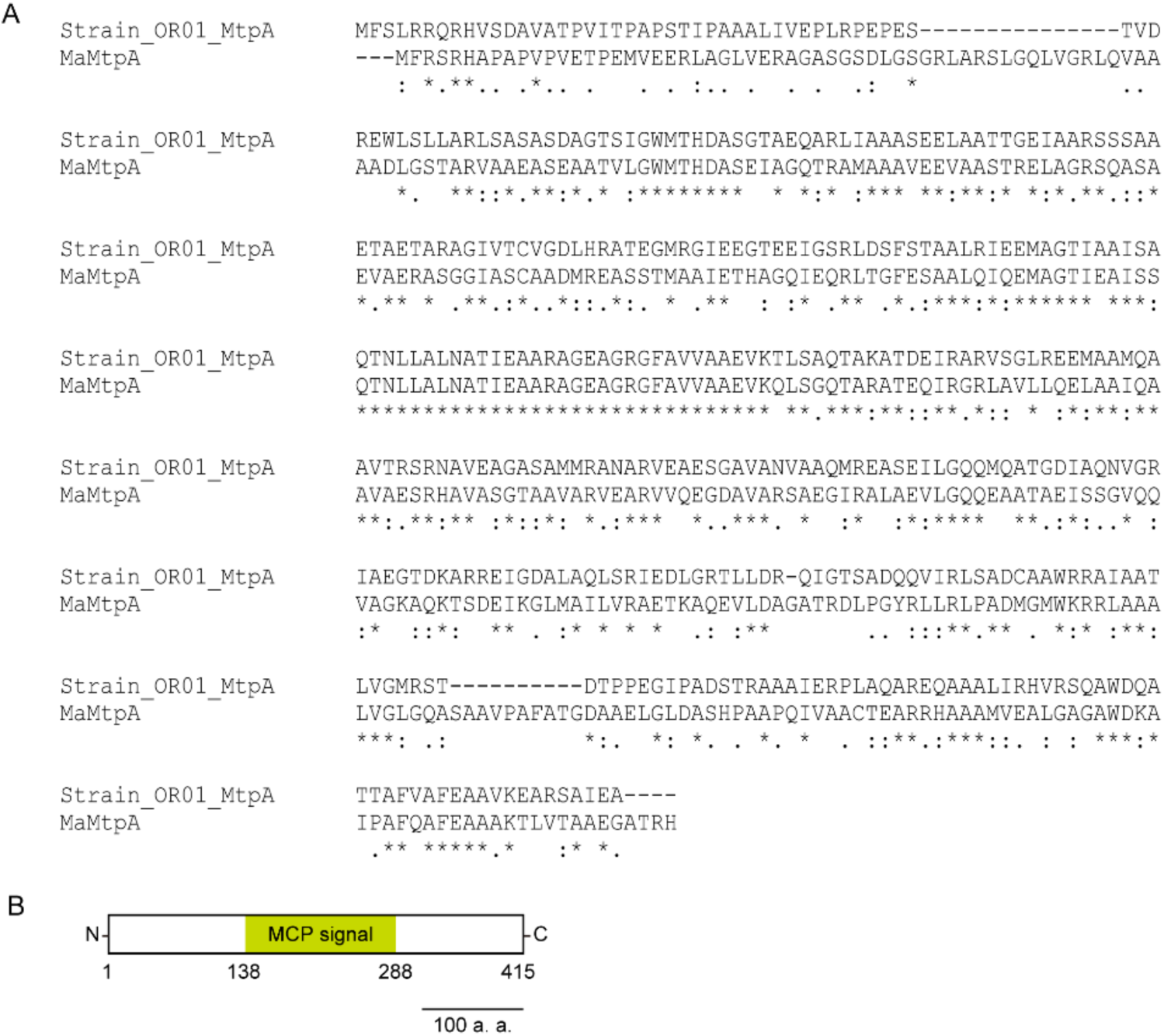
Comparison of the amino acid sequence of MtpA. **A** Alignment of MtpA in strain OR01 (Strain_OR01_MtpA) with MaMtpA in strain 22A. MtpA in strain OR01 contains a 1242-bp ORF encoding a 414-amino acid protein. **B** Schematic diagram of MtpA structure from strain OR01. The conserved motifs were analyzed by GenomeNet MOTIF Search (https://www.genome.jp/tools/motif/), as described previously(Tani et al., 2023). Methyl-accepting chemotaxis protein (MCP) signaling domain, demonstrated as MCP signal, is identified. Bar, 100 amino acids.

**Fig 4—figure supplement 2.**
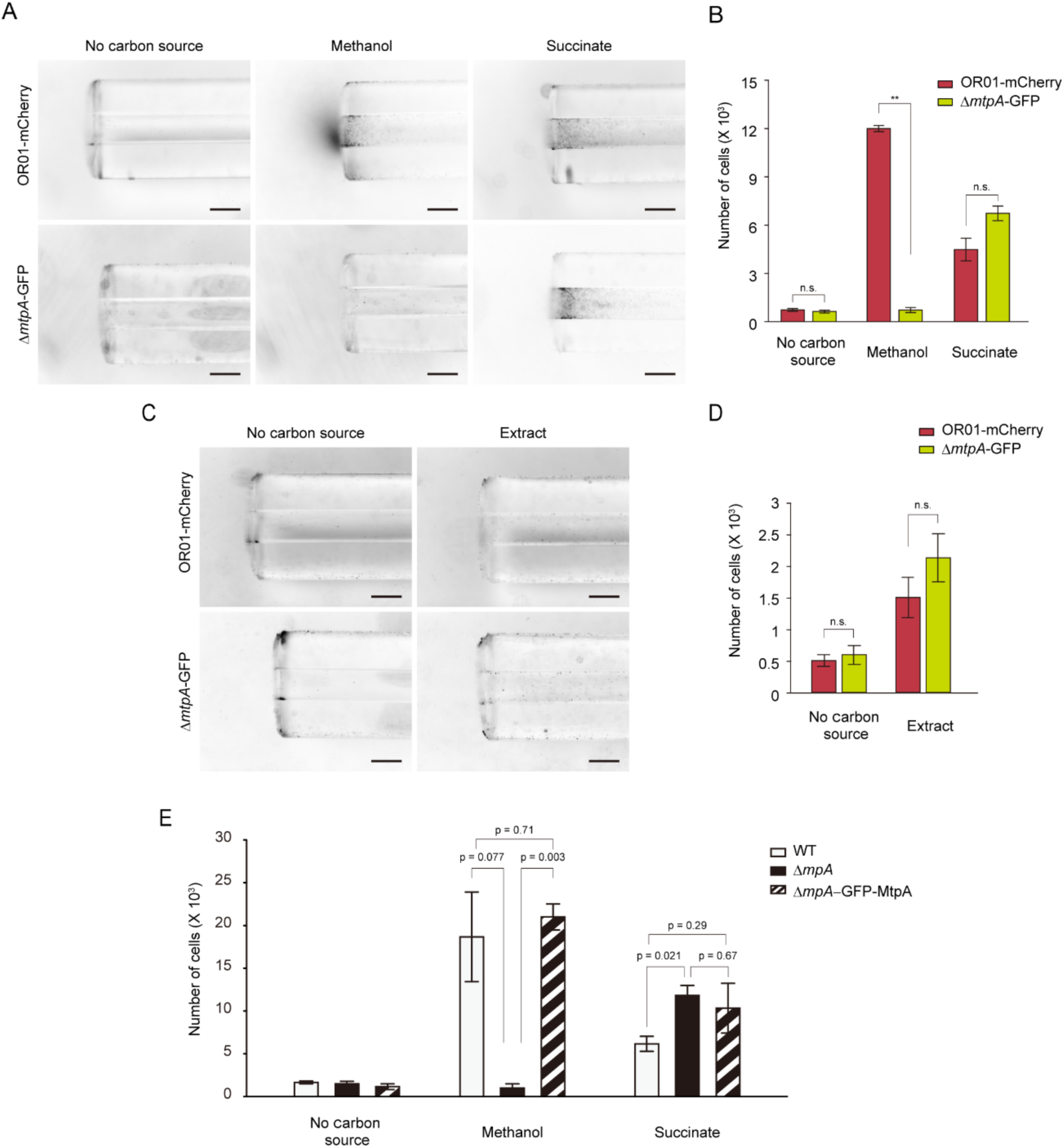
Deletion of *mtpA* gene led to significant reduction in methylotaxis. **A, C** Capillary assay for chemotactic response of strain OR01-mCherry and strain Δ*mtpA*-GFP. These strains were placed close to the capillaries that contain a hypho minimal medium supplemented with 0.05% methanol or 0.05% succinate (**A**) or 0.4% extract from red perilla leaves (**C**). Hypho minimal medium without any carbon source was used as a control. Images were taken by a stereo microscope. Bar, 200 µm. **B, D** FCM-based quantification of cell populations of strain OR01-mCherry and strain Δ*mtpA*-GFP in capillaries used in **A** and **C**, respectively. Values are indicated as the number of cells per capillary and are shown as mean ± s.e.m. of three independent measurements. Asterisks indicate level of statistical significance: * p < 0.05. Not significant: n.s. **E** Complementation analysis of methylotaxis using WT strain, strain Δ*mtpA* and strain Δ*mtpA*-GFP-MtpA in hypho minimal medium supplemented with 0.05% methanol or 0.05% succinate. Cells in each of the capillaries were spread onto hypho minimal agar plates supplemented with 0.5% methanol. After 2–3 days when colonies appeared, the cell number was investigated by FCM. Values are indicated as the number of cells per capillary and are shown as mean ± s.e.m. of three independent measurements. Statistical significance is shown as p values.

**Fig 5—figure supplement 1.**
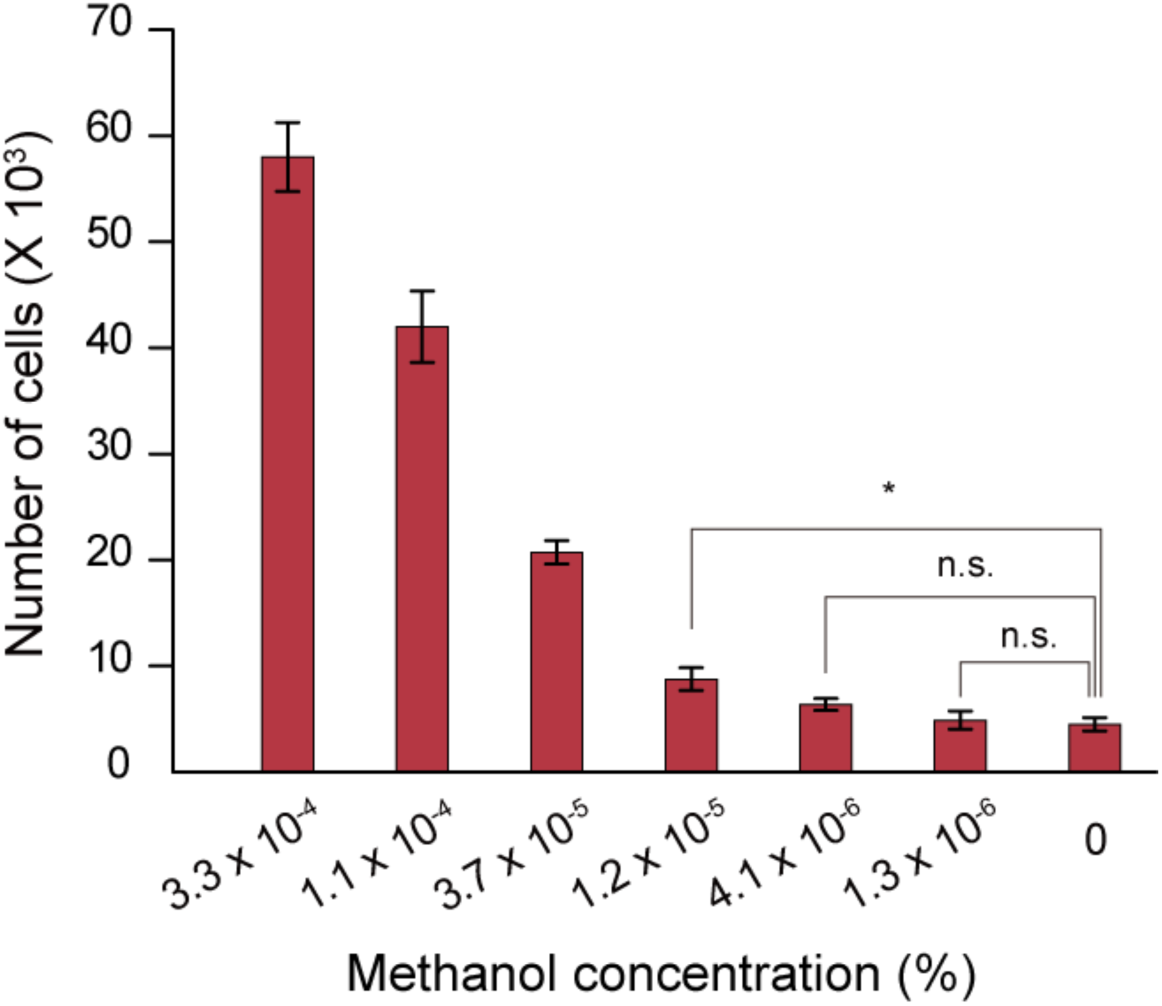
Capillary assay for chemotactic response of strain OR01-mCherry. Strain OR01-mCherry was placed close to the capillaries that contained a hypho minimal medium supplemented with methanol at the indicated concentration and incubated at 28°C for 24 hours. Cell populations of strain OR01-mCherry in capillaries were quantified by FCM. Values are indicated as the number of cells per capillary and are shown as mean ± s.e.m. of four independent measurements. Asterisks indicate level of statistical significance: * p < 0.05. Not significant: n.s.

## Videos

**Video 1. Z-stack videography of strain OR01-GFP on the red perilla leaf surface with a focus on stomata.**

Z-stack images of stomata with strain OR01-GFP taken by a confocal fluorescence microscope (see also Fig. 2C). Sixteen images were taken with 2 µm depth in the z-axis direction per 0.5 second.

**Video 2. Videography of strain OR01-GFP entering into the stomatal cavity.**

Time-lapse images of strain OR01-GFP entering into the stomata cavity (see also Fig. 2F). The video was taken by a confocal microscope and recorded for 77 seconds at 1x speed. Bar, 20 µm.

**Video 3. Motility analysis of wild-type cells.**

Videography of wild-type (WT) cells grown on hypho minimal medium containing 0.5% methanol and transferred to slide glasses for analysis (see also Figure 3—figure supplement 1C). Videos were taken at about 10 seconds. Bar, 20 µm.

**Video 4. Motility analysis of Δ*fliC-*triple cells.**

Videography of Δ*fliC-*triple cells grown on hypho minimal medium containing 0.5% methanol and transferred to slide glasses for analysis (see also Figure 3—figure supplement 1D). Videos were taken at about 10 seconds. Bar, 20 µm.

## Supplementary File

**Supplementary File 1A: Bacterial strains used in this study**

**Supplementary File 1B: Composition of hypho minimal medium**

**Supplementary File 1C: Plasmids used in this study**

**Supplementary File 1D: Oligonucleotide primers used in this study.**

## Notes

### Competing Interest Statement

The authors have declared no competing interest.

### Summary of Updates

Supplementary Fig. 5 (Fig 5-figure supplement 1) was updated and related descriptions in the main text were revised.

