## Supplementary files 1A to 1D for "Methanol chemoreceptor MtpA- and flagellin protein FliC-dependent methylotaxis determines the spaciotemporal colonization of PPFM in the phyllosphere"

**Supplementary File 1A: Bacterial strains used in this study**

| Strain | Genotype | Reference |
|---|---|---|
| ***Methylobacterium* sp. strain OR01** | | |
| OR01 | Wild-type | Mizuno *et al*. 2013(Mizuno, Yurimoto, Iguchi, Tani, & Sakai, 2013) |
| OR01-GFP | OR01::(pCM802-GFP, $Km^r$) | This study |
| OR01-mCherry | OR01::(pCM802-mCherry, $Km^r$) | This study |
| OR01-mCherry-g | OR01 locus 402468-403563::(pCM1684, $Km^r$) | This study |
| Δ*fliC1* | OR01Δ*fliC1* | This study |
| Δ*fliC2* | OR01Δ*fliC2* | This study |
| Δ*fliC*-double | OR01Δ*fliC1*Δ*fliC2* | This study |
| Δ*fliC*-triple | OR01Δ*fliC1*Δ*fliC2*Δ*fliC3* | This study |
| Δ*fliC*-triple-GFP | Δ*fliC1*Δ*fliC2*Δ*fliC3*::(pCM802-GFP, $Km^r$) | This study |
| Δ*mtpA*-GFP | Δ*mtpA*::(pCM802-GFP, $Km^r$) | This study |
| Δ*mtpA*-GFP-MtpA | Δ*mtpA*::(pCM802-GFP-mtpA, $Km^r$) | This study |
| ***M. aquaticum* strain 22A** | | |
| 22A | Wild-type | Tani *et al.* 2015(Tani, Ogura, Hayashi, & Kimbara, 2015) |
| 22A-mVenus-g | 22A locus 1717648-1719862::(pCM1685, $Km^r$) | This study |

**Supplementary File 1B: Composition of hypho minimal medium**

| Reagent | Final concentration (mg/L) |
|---|---|
| $K_2HPO_4$ | 2530 |
| $NaH_2PO_4$ | 2250 |
| $(NH_4)_2SO_4$ | 500 |
| $MgSO_4$ 7 $H_2O$ | 200 |
| EDTA 2Na | 12.74 |
| $ZnSO_4$ $7H_2O$ | 4.4 |
| $CaCl_2$ $2H_2O$ | 1.466 |
| $MnCl_2$ $4H_2O$ | 1.012 |
| $FeSO_4$ $7H_2O$ | 0.998 |
| $(NH_4)_6Mo_7O_{24}$ $4H_2O$ | 0.22 |
| $CuSO_4$ $5H_2O$ | 0.314 |
| $CoCl_2$ $6H_2O$ | 0.322 |

The following Vitamin mix was added to the medium, as necessary.

| Reagent | Final concentration (μg/L) |
|---|---|
| Ca-pantothenate | 400 |
| Inositol | 200 |
| Niacin (nicotinic acid) | 400 |
| p-Aminobenzonate | 200 |
| Pyridoxine HCl | 400 |
| Biotin | 2 |
| Thiamin HCl | 400 |

**Supplementary File 1C: Plasmids used in this study**

| Plasmid | Description | Reference |
|---|---|---|
| **Fluorescent markers** | | |
| pBluescript II SK(+) | Cloning vector, $Ap^r$ | Stratagene Inc. |
| pDCG-1 | Source of GFP, $Ap^r$ $Cm^r$ | Iguchi *et al.* 2013 (H. Iguchi, Sato, Yurimoto, & Sakai, 2013) |
| pMO149 | Source of mCherry, $Ap^r$ | Maeda *et al.* 2015(Maeda, Oku, & Sakai, 2015) |
| pAT02-V | Source of mVenus, $Ap^r$ | Tani *et al.* 2023(Tani et al., 2023) |
| pBS-$P_{mxaF}$ | pBluescript II SK(+) carrying the *mxaF* promoter of *Methylobacterium* sp. strain OR01 | This study |
| $P_{mxaF}$-GFP | pBS-$P_{mxaF}$ expressing *GFP* under the *mxaF* promoter of *Methylobacterium* sp. strain OR01 | This study |
| $P_{mxaF}$-mCherry | pBS-$P_{mxaF}$ expressing *mCherry* under the *mxaF* promoter of *Methylobacterium* sp. strain OR01 | This study |
| pCM80-Km | Expression vector carrying the *mxaF* promoter of *M. extorquens* strain AM1, $Km^r$ | Orita *et al*. 2014(Orita, Nishikawa, Nakamura, & Fukui, 2014) |
| pCM802-Km | pCM80-Km without carrying the *mxaF* promoter of *M. extorquens* strain AM1, $Km^r$ | This study |
| pCM802-GFP | pCM802-Km expressing *GFP* under the *mxaF* promoter of *Methylobacterium* sp. strain OR01 | This study |
| pCM802-mCherry | pCM802-Km expressing *mCherry* under the *mxaF* promoter of *Methylobacterium* sp. strain OR01 | This study |
| pCM802-GFP-mtpA | pCM802-Km expressing *GFP-mtpA* under the *mtpA* promoter, $Km^r$ | This study |
| pCM1682 | Allelic exchange vector with *katA*::$Km^r$ | Iguchi *et al.* 2018(Hiroyuki Iguchi et al., 2018) |
| pCM1684 | Allelic exchange vector with (genome region of strain OR01 at 402468-403563 locus)::$Km^r$, expressing *mCherry* under the *mxaF* promoter | This study |

| | | |
|---|---|---|
| pCM1685 | Allelic exchange vector with (genome region of strain 22A at 1717648-1719862 locus)::Km$^r$, expressing *Venus* under the *mxaF* promoter | This study |
| **Gene deletion** | | |
| pK18 mobsacB | Cloning vector, mob, sacB, Km$^r$ | Iguchi *et al.* 2013(H. Iguchi et al., 2013) |
| pK18 fliC1 | pK18 mobsacB harboring *fliC1* and the homologous fragments flanking *fliC1* | This study |
| pK18 ΔfliC1 | pK18 mobsacB harboring the homologous fragments flanking *fliC1* | This study |
| pK18 fliC2 | pK18 mobsacB harboring *fliC2* and the homologous fragments flanking *fliC2* | This study |
| pK18 ΔfliC2 | pK18 mobsacB harboring the homologous fragments flanking *fliC2* | This study |
| pK18 ΔfliC3 | pK18 mobsacB harboring the homologous fragments flanking *fliC3* | This study |
| pK18 mtpA | pK18 mobsacB harboring *matpA* and the homologous fragments flanking *mtpA* | This study |
| pK18 ΔmtpA | pK18 mobsacB harboring the homologous fragments flanking *mtpA* | This study |

**Supplementary File 1D: Oligonucleotide primers used in this study**

| Primer names | Sequence (5'-3') |
|---|---|
| **Fluorescent markers** | |
| pCM80km-Re1800 | GCGGTAATACGGTTATCCAC |
| pCM80km-Fw2471 | CGCCAAGCTTGCATGCCTGC |
| PmxaF-Fw-KpnI | GGGGTACCACAGGTCGCCAGCGCCAGAA |
| PmxaF-Rv-HindIII | CCCAAGCTTCCTGCGTCTCCTCGCCGGAC |
| GFP-Fw-HindIII | CCCAAGCTTATGAGTAAAGGAGAAGAACT |
| GFP-Rv-PstI | AACTGCAGTTATTTGTAGAGCTCATCCA |
| mCherry-Fw-HindIII | CCCAAGCTTATGGTGAGCAAGGGCGAG |
| mCherry-Rv-BamHI | AAGGATCCAGGACTTGTACAGCTCG |
| 1vector-Fw | TCATCTCAACATTATTTGAATACAGGGGCATCGACG |
| 1vector-Rv | CAGAACCGGCCCACGCGA |
| 3vector-Fw | CGCACAAGATGCCATGTATGG |
| 3vector-Rv | CGCGAGGCGATATCGTCCATTC |
| OR01-2insertB-Fw | CGTGGGCCGGTTCTGCAACATCGTGCTCTACGCC |
| OR01-2insertB-Rv | ATGGCATCTTGTGCGGGTGTTGAACGAGAAGCCG |
| OR01-4insertB-Fw | CGATATCGCCTCGCGCTGCTCGGCATGGTGCTG |
| OR01-4insertB-Rv | ATAATGTTGAGATGAGCCGGTGCCTCAGGATCAG |
| PmxaF-KpnI-Fw | CGAGCTCCCGGGTACACAGGTCGCCAGCGCCAGAAA |
| mCherry-KpnI-Rv | TGCATGCCATGGTACCTTGTACAGCTCGTCCATGCCGCC |
| 22A-2insertC-Fw | CGTGGGCCGGTTCTGCGCACATCTTCCGCCGTAC |
| 22A-2insertC-Rv | ATGGCATCTTGTGCCGGAAGATGTGCGCCACCTCC |
| 22A-4insertC-Fw | CGATATCGCCTCGCGGACGGCGGAAAGCGTGGAG |
| 22A-4insertC-Rv | ATAATGTTGAGATGACGATATGCTGGTCGACATGC |
| EcoRI-PmxaF-Fw | TTGGTTGTAACACTGAATTCGCCGATGTCACCGTGCTG |
| GFP-EcoRI-Rv | CTTAAGCTCGAGGGCCCATGTAACTTGTACAGCTCGTCCA TGCCG |
| NB_PmtpA_Fw | AGGTCGACTCTAGAGGCCGAGACGGAGGATGCTC |
| NB_PmtpA+GFP_Rv | CTTTACTCATGCGGGACACTCCGATATCCG |
| NB_PmtpA+GFP_Fw | CCCGCATGAGTAAAGGAGAAGAACTTTTCACTGGAG |
| NB_GFP+mtpA_Rv | CTAAACATTTTGTAGAGCTCATCCATGCCATGTG |
| NB_GFP+mtpA_Fw | TCTACAAAATGTTTAGTTTGCGTCGGCAGC |
| NB_mtpA_Rv | CAGTGAATTCGAGCTTCAGGCCTCGATCGCCG |
| **Gene deletion** | |
| fliC1_up_Fw | GGGGATCCTCTAGAGTGGCGAGATCCATGGCTTC |
| fliC1_down_Rv | CCAGTGCCAAGCTTGTCCAGATCGGAGGTGCATC |
| Inv_fliC1_Fw | TCGCCGATCTGATTCAGGC |
| Inv_fliC1_Rv | GCGGCGTTGTTCGTAAACAG |
| fliC2_up_Fw | GGGGATCCTCTAGAGGATCTCTACGTCCGTCACACCG |
| fliC2_down_Rv | TGCCAAGCTTGCATGCCAATCCCCTGCTCAATGCAG |
| Inv_fliC2_Fw | GTTTGCATGAAAAGCGTCGG |
| Inv_fliC2_Rv | GAGCAGGTTCTGGCGTGTC |
| fliC3_up_Fw | GGGGATCCTCTAGAGTTGTCGTAATGCCATTTCGAG |
| fliC3_up_Rv | CGTCCGGTCAATGGTATCCTGCAGGGAGAGCAG |

| | |
|---|---|
| fliC3_down_Fw | ACCATTGACCGGACGGACGAGATG |
| fliC3_down_Rv | CCAGTGCCAAGCTTGGGCCTCAAAGATCCGGATCGCG |
| mtpA_up_Fw | TCGAGCTCGGTACCCCCCACCACGGAGTTCGAG |
| mtpA_ down_Rv | CTCTAGAGGATCCCCAGATCGAGTTCTGGCTGCTG |
| Inv_mtpA_Fw | CGTCGCGGGTCGTCTTAC |
| Inv_mtpA_down | CTCTAGAGGATCCCCAGATCGAGTTCTGGCTGCTG |
